# A potent synthetic nanobody targets RBD and protects mice from SARS-CoV-2 infection

**DOI:** 10.1101/2020.06.09.143438

**Authors:** Tingting Li, Hongmin Cai, Hebang Yao, Bingjie Zhou, Ning Zhang, Yuhuan Gong, Yapei Zhao, Quan Shen, Wenming Qin, Cedric A.J. Hutter, Yanling Lai, Shu-Ming Kuo, Juan Bao, Jiaming Lan, Markus A. Seeger, Gary Wong, Yuhai Bi, Dimitri Lavillette, Dianfan Li

## Abstract

SARS-CoV-2, the causative agent of COVID-19^1^, recognizes host cells by attaching its receptor-binding domain (RBD) to the host receptor ACE2^2–7^. Neutralizing antibodies that block RBD-ACE2 interaction have been a major focus for therapeutic development^8–18^. Llama-derived single-domain antibodies (nanobodies, ∼15 kDa) offer advantages including ease of production and possibility for direct delivery to the lungs by nebulization^19^, which are attractive features for bio-drugs against the global respiratory disease. Here, we generated 99 synthetic nanobodies (sybodies) by *in vitro* selection using three libraries. The best sybody, MR3 bound to RBD with high affinity (*K*_D_ = 1.0 nM) and showed high neutralization activity against SARS-CoV-2 pseudoviruses (IC_50_ = 0.40 μg mL^−1^). Structural, biochemical, and biological characterization of sybodies suggest a common neutralizing mechanism, in which the RBD-ACE2 interaction is competitively inhibited by sybodies. Various forms of sybodies with improved potency were generated by structure-based design, biparatopic construction, and divalent engineering. Among these, a divalent MR3 conjugated with the albumin-binding domain for prolonged half-life displayed highest potency (IC_50_ = 12 ng mL^−1^) and protected mice from live SARS-CoV-2 challenge. Our results pave the way to the development of therapeutic nanobodies against COVID-19 and present a strategy for rapid responses for future outbreaks.

## INTRODUCTION

The coronavirus disease that emerged in early December 2019 (COVID-19)^1^ poses a global health and economic crisis^20^. The causative agent, SARS-CoV-2, uses its Spike protein (S) to recognize receptors on host cells, an initial step for viral infection^2,3,21,22^. Key to this virus-host interaction is the binding between the S receptor-binding domain (RBD) and the host ACE2 protein^4–7^. Therefore, the RBD has been a primary target for neutralizing antibodies^8–13,23^ to block ACE2-binding.

Llama-derived nanobodies are generally more heat stable, easier and less expensive for production, and more amenable to protein engineering compared to conventional antibodies^24^. As single-chain antibodies, nanobody libraries are less complex to construct and screen, enabling *in vitro* selection of high-affinity binders in relative short time, typically 2-4 weeks^25–30^. Recently, several nanobody therapeutics, including the caplacizumab approved by the US Food and Drug Administration, have been developed for a variety of immune diseases^31^. Of relevance to SARS-CoV-2, nanobodies can survive nebulization and an inhaler nanobody drug (ALX-0171) has gone into clinical trials for the treatment of the Respiratory Syncytial Virus^31^. Recent weeks have witnessed the generation of nanobodies that neutralize SARS-CoV-2 from several independent groups^14–18^. However, the *in vivo* efficacy of such nanobodies remains to be investigated.

Here, we report our efforts in selection and engineering synthetic nanobodies (sybodies)^26^ that are highly potent against SARS-CoV-2, using biochemical and structural approaches. For the first time, we demonstrate that nanobodies can protect mice from live SARS-CoV-2 infection. Our results form a preliminary basis for the development of nanobody therapeutics for COVID-19.

## RESULTS AND DISCUSSION

### Generation of high-affinity neutralizing sybodies against SARS-CoV-2

SARS-CoV-2 S-RBD binders were selected by performing one round of ribosome display using three high-diversity libraries (Concave, Loop, and Convex)^26,27^, and three rounds of phage display using the RBD as the bait under increasingly stringent conditions. Subsequent ELISA (**Extended Data Fig. 1**) identified 80, 77, and 90 positive clones, corresponding to 62, 19, and 18 unique binders from the Concave, Loop, and Convex library, respectively (**Extended Data Table 1**). Eighty sequencing ‘first-comers’ of the 99 sybodies were further screened by a convenient fluorescence-detector size exclusion chromatography (FSEC) assay using crude extract from sybody-expressing clones. This identified 28 (36%) sybodies, including 9 Concave (21%), 9 Loop (50%), and 10 Convex (56%) binders that caused earlier retention of the fluorescein-labeled RBD (**Extended Data Fig. 2A, Extended Data Table 1**).

The same 80 sybodies were also screened for neutralization activity against retroviral pseudotypes harboring the SARS-CoV-2 S protein. Using 50% neutralization at 1 μM concentration as a cut-off, 11 Concave (26%), 13 Loop (68%), and 10 Convex (56%) sybodies were identified as positive (**Extended Data Fig. 3A**). The high positive rates suggest high efficiency of the *in vitro* selection platform. Of note, none of the sybodies showed noticeable neutralization activities for the closely related SARS-CoV pseudovirus (**Extended Data Fig. 3B**), indicating high specificity.

Six FSEC-positive neutralizing sybodies, namely SR4 (1), MR3 (31), MR4 (9), MR17 (1), LR1 (31), and LR5 (19) (S, M, L refers to Concave, Loop, and Convex sybodies respectively; brackets indicate ELISA redundancy), were characterized in more detail as follows. They could be purified from *Escherichia coli* with high yield (**Fig. 1A**), formed complexes with RBD on gel filtration (**Extended Data Fig. 2B**), displayed ultra-high thermostability (**Fig. 1A, Extended Data Fig. 2C**) as originally designed ^26^, and bound to the RBD with relatively high affinity (**Fig. 1A, Extended Data Fig. 4)**, with *K*_D_ ranging from 83.7 nM (MR17) to 1.0 nM (MR3). Consistent with its highest affinity, MR3 showed the slowest off-rate (2.3 × 10^−4^ s^−1^). Using neutralization assays, we determined IC_50_ of the six sybodies (**Fig. 1B**). MR3 was the most potent (IC_50_ of 0.40 μg mL^−1^), indicating a largely consistent trend between neutralization potency and binding kinetics (affinity and off-rate).

**Fig. 1.**
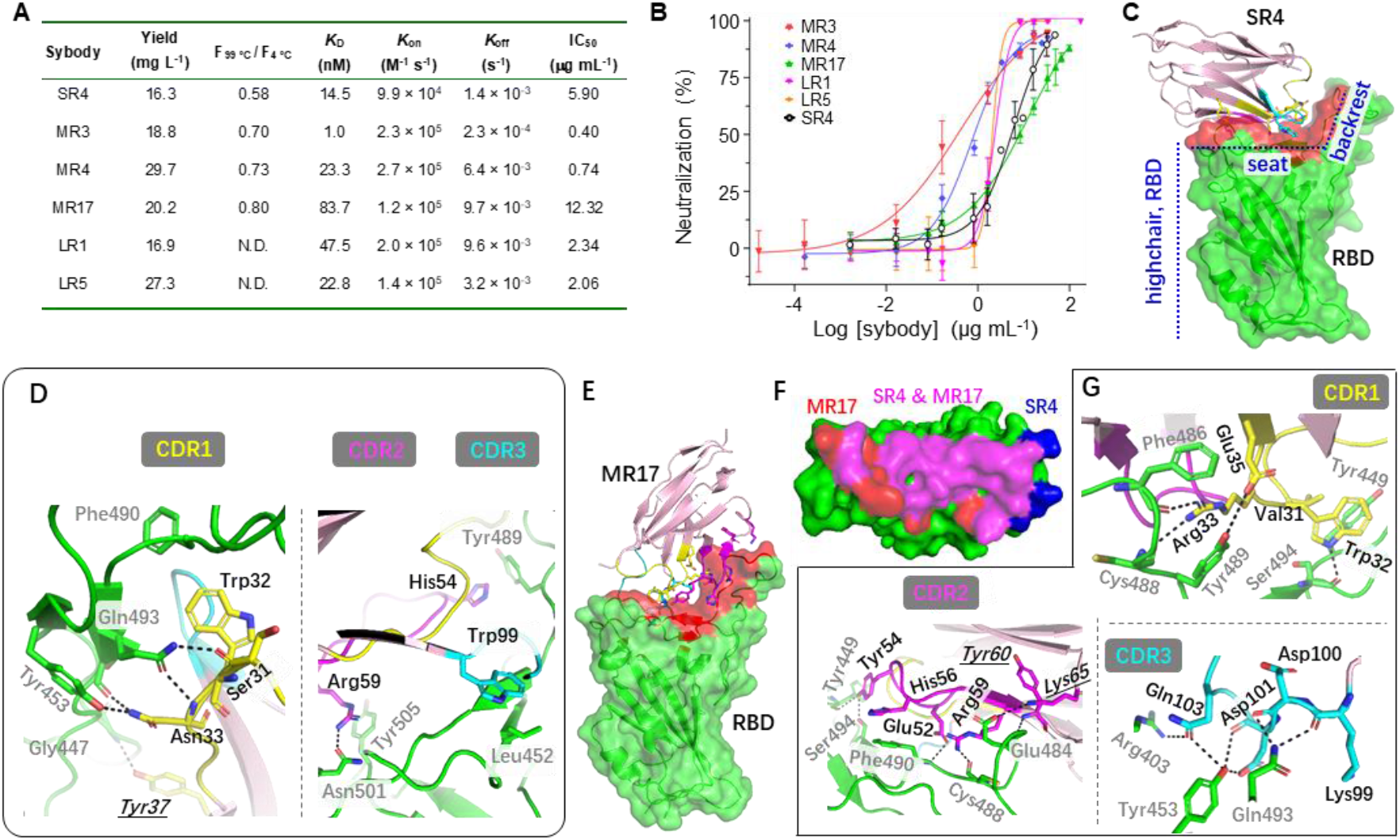
Biochemical and structural characterization of neutralizing sybodies. (**A**) Summary of the characterization. Yield refers to purification from 1 L of culture. Fractional fluorescence (F) indicates remaining gel filtration peak intensity of sybodies after heating at 99 °C for 20 min. N. D., not determined. (**B**) Neutralization assay. SARS-CoV-2 pseudoviruses were pre-incubated with different concentration of sybodies before infection of VeroE6-hACE2 cells. The rate of infection was measured by fluorescence-activated cell sorting (FACS). IC_50_ was obtained by Sigmoidal fitting of the percentage of neutralization. Data are from three independent experiments. (**C**) The overall structure of SR4 (pink cartoon) in complex with RBD (green surface) which resembles a short backrest high chair. The binding surface is highlighted red. (**D**) SR4 CDR1 (yellow), CDR2 (magenta), and CDR3 (cyan) all contributed to the binding. Underlining italics label the framework residue Tyr37. (**E**) The overall structure of the MR17 (pink cartoon) in complex with RBD (green surface). (**F**) The overlap (magenta) between the SR4-(blue) and MR17-(red) interacting surfaces on RBD. (**G**) All three CDRs contributed to the binding with RBD (green). Underlining italics label the framework residues Lys65 and Tyr60. Dashed lines indicate H-bonding or salt-bridges between atoms that are <4.0 Å apart. Black texts label sybody residues and grey texts label RBD residues.

### Structure of sybody-RBD complexes

To gain mechanistic insights into neutralization, we performed crystallization trials for several RBD-sybody complexes and obtained crystals for four. Crystals of SR4- and MR17-RBD diffracted to 2.15 Å and 2.77 Å resolution respectively and allowed structure determination (**Extended Data Table 2**). Crystals for MR3- and MR4-RBD did not diffract beyond 8.0 Å despite our optimization efforts.

The RBD structure resembles a short backrest high chair and SR4 binds to both the ‘seat’ and ‘backrest’ (**Fig. 1C**) with a surface area^32^ of 727.37 Å^2^ with modest electrostatic complementarity (**Extended Data Fig. 5A**). Of note, SR4 binds sideways, as intended by design of the Concave sybody library^26^. All three CDRs contributed to the binding through hydrophobic interactions and H-bonding that involves both side chains and main chains (**Fig. 1D**). In addition, Tyr37, a framework residue, also participated binding by forming an H-bond with the RBD Gly447 backbone.

MR17 also binds to the RBD at the ‘seat’ and ‘backrest’ regions but approaches the RBD at an almost perfect opposite direction of SR4 (**Fig. 1C, 1E**), indicating divergent binding mode for these sybodies. The binding of MR17 to the RBD occurred on an 853.94 Å^2^ surface area with noticeable electrostatic complementarity (**Extended Data Fig. 5B**). Interestingly, this surface was largely shared with the SR4 binding surface (**Fig. 1F**). The interactions between MR17 and the RBD were mainly mediated by H-bonding. Apart from the three CDRs, two framework residues, Lys65 and Tyr60, interacted with the same RBD residue Glu484, via a salt bridge with its side chain, and an H-bond with its main chain (**Fig. 1G**).

### Molecular mechanism for neutralization

Structure alignment of SR4-, MR17- and ACE2-RBD^4^ showed that both sybodies engage with RBD at the receptor-binding motif (RBM) (**Fig. 2A, 2B)**. Superposing SR4 and MR17 to the S trimer showed both sybodies could bind to the ‘up’ conformation^2^ of RBD with no steric clashes (**Fig. 2C, 2D**), and to the ‘down’ conformation with only minor clashes (**Extended Data Fig. 6**) owing to their minute sizes. Consistent with the structure observation, both SR4 and MR17 inhibited the binding of ACE2 to RBD, as revealed by bio-layer interferometry (BLI) assays (**Fig. 2E, 2F**).

**Fig. 2.**
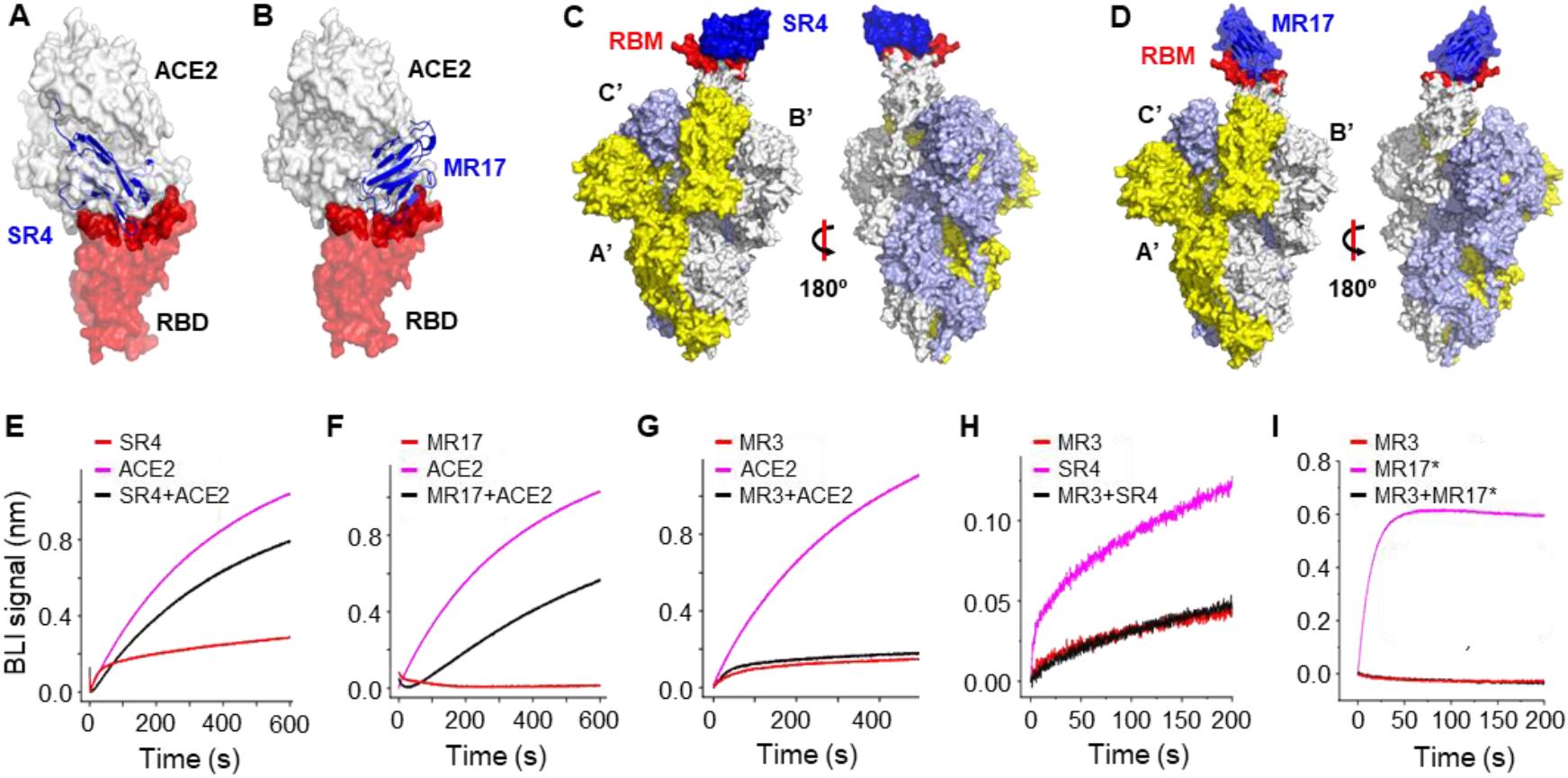
Molecular basis for neutralization. (**A**,**B**) Alignment of the SR4-(**A**) or MR17-(**B**) RBD to the ACE2-RBD structure (PDB ID 6M0J)^4^ reveals that SR4/MR17 (blue) binds RBD (red) at the motif (dark red) where ACE2 (white) also binds at. (**C**,**D**) Alignment of the SR4-RBD (**C**) and MR17-RBD (**D**) to the ‘up’ conformation of the RBD from the cryo-EM structure of the trimer S (PDB ID 6VYB)^2^. A’/B’/C’ label three subunits. RBM (red) marks the ACE2-binding motif. (**E-G**) Competitive binding for the RBD between sybody and ACE2. A sensor coated with streptavidin was saturated with 2 μg mL^−1^ of biotinylated RBD. The sensor was then soaked in 200 nM of indicated sybody before further soaked in sybody-containing buffer with (black) or without (red) 25 nM of ACE2 for BLI signal recording. As a control, the ACE2-RBD interaction was monitored in the absence of sybodies (magenta). (**H-I**) Competitive BLI assay for the RBD between sybody pairs. A sensor with immobilized RBD was soaked in 200 nM of MR3 before further soaked in MR3-containing buffer with (black) or without SR4/MR17 (red). As a control, the SR4- and MR17-RBD interaction were monitored in the absence of MR3 (magenta). In (**I**), MR17* indicates a MR17 mutant (see below). Panels **E-I** share the same Y-axis title.

To probe the epitope for MR3 without a structure, competitive BLI binding assays were carried out. The results showed that MR3 could block ACE2 (**Fig. 2G**), and SR4 and MR17 (**Fig. 2H, 2I**), suggesting it also binds to at least part of the RBM, although the possibility of allosteric inhibition remains to be investigated. Taken together, SR4 and MR17, and probably MR3, neutralize SARS-CoV-2 by competitively blocking the ACE2-RBD binding.

### Sybody engineering increased affinity and neutralizing activity

Increasing valency is a common technique to enhance potency for nanobodies^18,31^. To this end, we engineered three types of divalent sybodies, including the biparatopic fusion of two different sybodies, the Fc-fusion and direct fusion of the same sybody.

For biparatopic fusion, we first identified two sybodies, namely LR1 and LR5 (**Fig. 3A, 3B**), that could bind RBD in addition to MR3 using the BLI assay. As LR5 showed higher affinity and neutralization activity than LR1 (**Fig. 1A**), we fused this non-competing sybody to the N-terminal of MR3 with various length of GS linkers ranging from 13 to 34 amino acids (**Extended Data Table S1**). Interestingly, the linker length had little effect on neutralization activity and these biparatopic LR5-MR3 sybodies were more potent than either sybodies alone (**Fig. 1A**) with an IC_50_ of 0.11 μg mL^−1^ (**Fig. 3C**). LR5-MR3 may be more tolerant to escape mutants^34–37^ owing to its ability to recognize two distinct epitopes.

**Fig. 3.**
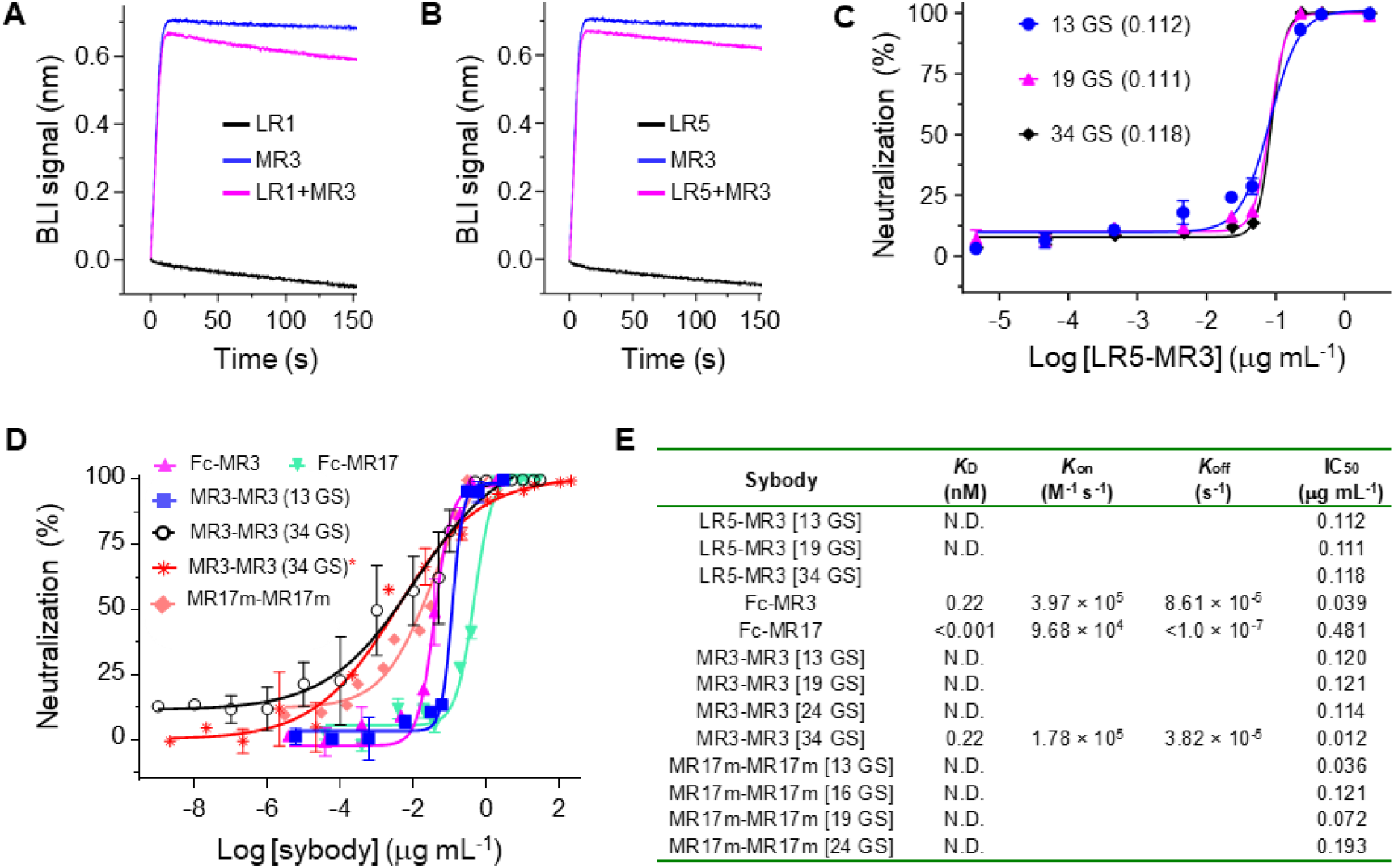
Divalent engineering increased affinity and neutralizing activity. (**A**,**B**) Identification of two non-competing pairs, LR1/MR3 (**A**) and LR5/MR3 (**B**), for biparatopic constructs. For BLI assays, sensors coated with RBD were soaked in 200 nM of LR1 or LR5 before further soaked in LR1- or LR5-containing buffer with (magenta) or without (black) 100 nM of MR3. The MR3-RBD interaction profile was obtained in the absence of LR1 or LR5 (blue). (**C**) Neutralization assay of the biparatopic sybody LR5-MR3 with a GS linker of various length as indicated. Brackets indicate IC_50_ values in μg mL^−1^. (**D**) Neutralization assays of divalent sybodies. The original SARS-CoV-2 was used for all assays except that the D614G mutant^33^ was additionally tested for MR3-MR3 (red asterisk). (**E**) Summary of binding kinetics and neutralizing activities of the divalent sybodies. N.D., not determined.

For Fc-fusion, both MR3 and MR17 were attached to the dimeric human IgG Fc. This decreased IC_50_ by 10 folds for Fc-MR3 (39 ng mL^−1^) and 25 folds Fc-MR17 (0.48 μg mL^−1^), respectively (**Fig. 3D, 3E**). Consistently, the Fc fusion increased the apparent binding affinity for both sybodies, with a *K*_D_ of 0.22 nM for Fc-MR3 and less than 1 pM for Fc-MR17 (**Extended Data Fig. 4H, 4I**). Note, however, Fc-MR17 did not gain as much neutralization potency as for the apparent binding affinity.

For direct fusion, MR3 and a rationally designed MR17 mutant (MR17m, **Extended Data Fig. 7**) that showed comparable IC_50_ with MR3 by a single mutation K99Y (0.50 μg mL^−1^, **Extended Data Fig. 7G**) were individually linked together via GS linkers with variable length ranging from 13 to 34 amino acids (**Extended Data Table 1**). The optimal construct for MR17m-MR17m had the shortest linker (13-GS) (**Fig. 3D, 3E**). By contrast, optimal neutralization activity was observed with the longest linker (34-GS) for MR3-MR3 (**Fig. 3D, 3E**). Again, MR3-MR3 was superior compared to MR17m-MR17m, showing a 2-fold higher neutralization activity with an IC_50_ of 12 ng mL^−1^ (**Fig. 3E**). Compared to the monovalent MR3 (IC_50_ of 0.40 μg mL^−1^), the divalent engineering increased the potency by over 30 folds. Notably, MR3-MR3 showed similar activity to inhibit pseudotypes harboring the original SARS-CoV-2 S or the current dominant and more infectious mutant D614G S (ref. ^33^) (**Fig. 3D**).

### Divalent MR3 protects mice from COVID-19

The most potent divalent sybody (MR3-MR3) was chosen to investigate the potential of nanobodies to protect mice from SARS-CoV-2 infection. Nanobodies have very short serum half-lives of several minutes due to their minute size^38^. To circumvent this, we fused MR3-MR3 to the N-terminus of an albumin-binding domain (ABD)^39^ which has been known to extend the circulating half-life of its fusion partners by increase in size and preventing intracellular degradation^31^. Conveniently, we expressed MR3-MR3-ABD in *Pichia pastoris*, which is the preferred host to express nanobody therapeutics owing to its robustness and its endotoxin-free production. Small-scale expression of MR3-MR3-ABD showed a secretion level of ∼250 mg L^−1^ with an apparent purity of >80% without purification (**Fig. 4A**). Note, this experiment was carried out using a shaker which gave cell density of OD_600_ of 16. Given its ability to grow to OD_600_ of 500 without compromising yield, the expression level of MR3-MR3-ABD may reach 7.5 g L^−1^ in fermenters. The potential for simple and high-yield production is especially attractive for the pandemic at a global scale.

**Fig. 4.**
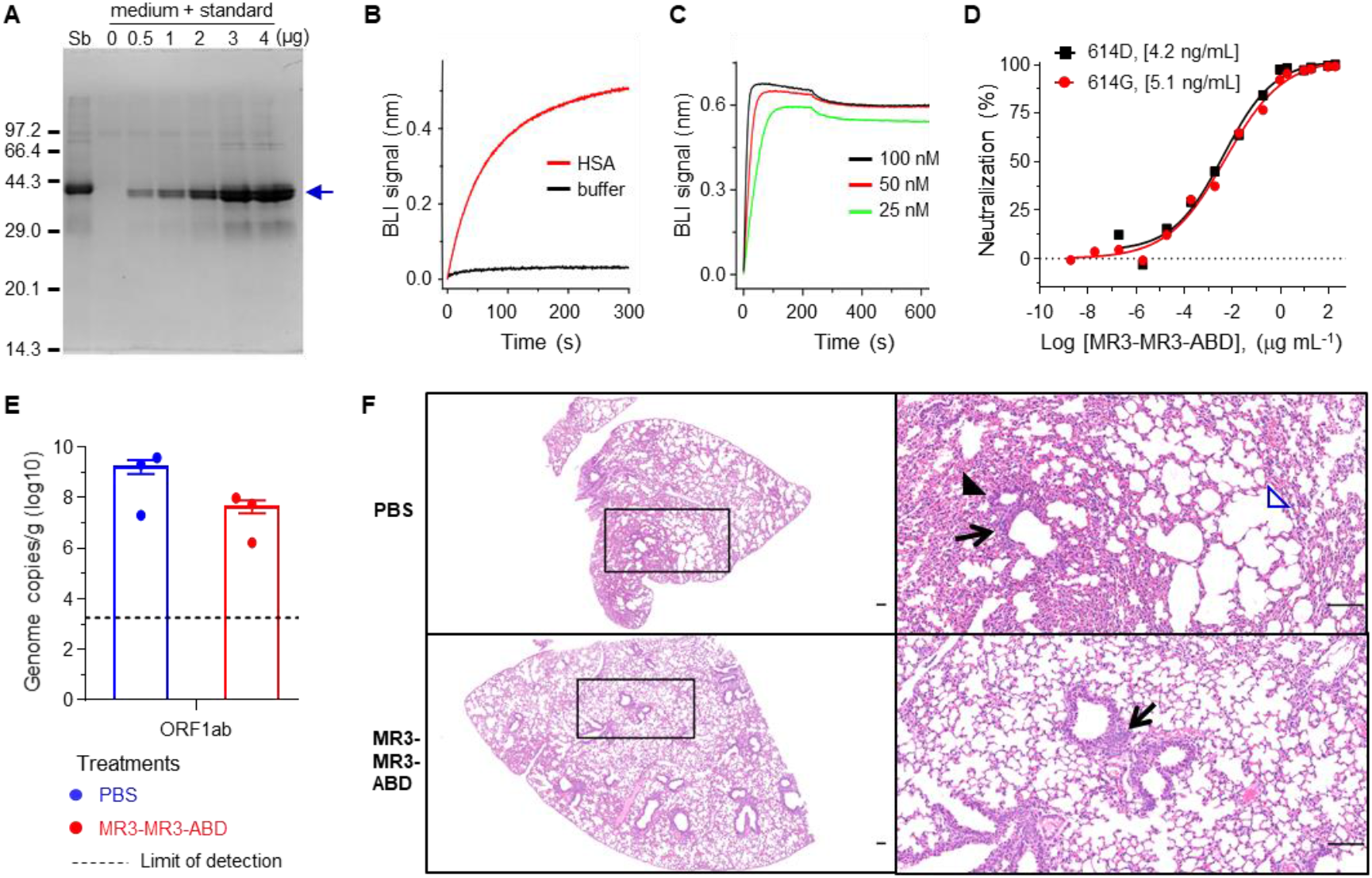
The Divalent MR3 sybody protects mice from live SARS-CoV-2 challenge. (**A**) Coomassie Blue staining of SDS-PAGE for MR3-MR3-ABD (arrow) expression in pichia. Based on the standards, the yield (Lane Sb) was semi-quantified as 0.25 g L^−1^. (**B**,**C**) BLI binding assays show that MR3-MR3-ABD bind to human serum albumin (HSA) (**B**) and RBD (**C**). RBD-coated sensors were incubated with 200 nM of HSA (**B**) or indicated concentrations of MR3-MR3-ABD (**C**) for single monitoring. (**D**) Neutralization assay of MR3-MR3-ABD. Data are from one representative experiment of two independent experiments. (**E**) Lung viral loads as determined by PCR from infected mice at 3 dpi. (**F**) Histopathology of lungs from infected mice at 3 dpi. The arrows denote inflammatory cell infiltration. A black triangle indicates typical red blood cell exudation, and a blue triangle indicates typical compensatory expansion of the alveolar cavity. The left panels denote an overview of the lung at 10x magnification. The right panels denote the expanded view of the black boxes in the left panels, at 100x magnification. Bars = 100 μm.

Importantly, MR3-MR3-ABD could bind to the human albumin (**Fig. 4B**) while retaining its ability to bind RBD (**Fig. 4C**) and to neutralize SARS-CoV-2 pseudotypes harboring either past (614D) and current (614G)^33^ SARS-CoV-2 S (**Fig. 4D**). As designed, a serum virus neutralization assay showed that the addition of the albumin binding domain to the divalent MR3 (MR3-MR3-ABD) extended its *in vivo* stability, displaying neutralization activity up to 24 h post injection contrary to the other forms (**Extended Data Fig. 8A**). The body weight measures, and the microscopic histopathology analysis did not reveal any toxicity for the nanobodies for 6 days (**Extended Data Fig. 8B, 8C**).

To test the *in vivo* antiviral efficacy of MR3-MR3-ABD, C57BL/6J female mice, aged 6-8 weeks old, were first sensitized to SARS-CoV-2 infection using an adenovirus expressing the human ACE2 receptor^40^ at 5 days before challenge. Mice were infected via the intranasal route with 5 x 10^6^ median tissue culture infectious dose (TCID_50_) of SARS-CoV-2, and then administered a single dose of 25 mg kg^−1^ MR3-MR3-ABD via the intraperitoneal route at 12 h after virus challenge. A control C57BL/6J group were given PBS as a mock treatment. Compared to the control group, the lung viral titers of the sybody group was 50-fold lower than the PBS group, when assessed at 3 dpi (**Fig. 4E**). This efficacy is similar to the existing human monoclonal antibody CB6 (2 injections, 50-fold, in rhesus macaques)^41^ and better than that for 1B07 (1 injection, 10-fold, mice)^42^ when compared under similar sampling points.

Histopathological examination revealed that the infected mice in the PBS-treated group displayed moderate bronchopneumonia lesions, with a large number of inflammatory cell infiltrations around the bronchioles and terminal bronchioles. The alveolar walls were thickened, a large number of inflammatory cells were exuded in the interstitium, accompanied by red blood cell exudation. In addition, part of the alveolar cavity showed compensatory expansion (**Fig. 4F**). In contrast, the lungs of sybody-treated mice showed normal alveolar wall structures, and only displayed mild bronchopneumonia, with a small amount of inflammatory cell infiltration around the bronchioles (**Fig. 4F**). Taken together, the significant reduction of the lung viral load and the severity of lung damage demonstrated the *in vivo* efficacy of the MR3-MR3-ABD against authentic SARS-CoV-2 infection.

In summary, the *in vitro* platform was efficient in generating neutralizing sybodies (the selection process took 2 weeks). Structural and biochemical studies suggested an antagonistic mechanism to block the ACE2-RBD interaction. Protein engineering yielded various forms of sybody with higher affinity, neutralization activity, and *in vivo* stability. Using the most potent construct, we have in the first time demonstrated that nanobodies can provide post-exposure protection of mice from SARS-CoV-2 infection. Our results should encourage development of nanobody therapeutics to fight COVID-19 or future viral outbreaks.

## ACKNOWLEDGMENTS

We thank the staff members of the Large-scale Protein Preparation System for equipment maintenance and management, and staff scientists at the SSRF-BL19U1 beamline at National Facility for Protein Science (Shanghai) for technical support and assistance. We thank Dr. Zhipu Luo at Soochow University (China) for helpful discussions regarding data processing. This work has been supported by the Strategic Priority Research Program of CAS (XDB37020204, D.Li; XDB29010102 & XDA19090118, Y.B.), Key Program of CAS Frontier Science (QYZDB-SSW-SMC037, D.Li), CAS Facility-based Open Research Program, the National Natural Science Foundation of China (31870726, D.Li; 31870153, D.La.; 32041010, Y.B.), the One Belt and One Road major project for infectious diseases (2018ZX10101004-003, J.L., G.W.), National Key R&D Program of China (2020YFC0845900, D.La.), CAS president’s international fellowship initiative (2020VBA0023, D.La.), Natural Science Foundation of Shanghai (20ZR1463900, G.W.), and Shanghai Municipal Science and Technology Major Project (20431900402, D.La.). Y.B. is supported by the NSFC Outstanding Young Scholars (31822055) and Youth Innovation Promotion Association of CAS (2017122). G.W. is supported by a G4 grant from IP, FMX and CAS.

## AUTHOR CONTRIBUTIONS

T.L., H.C., and H.Y. selected sybodies under the supervision of C.A.J.H. and M.A.S.. T.L., H.C., and H.Y. purified and crystalized protein complexes with assistance from Y.L.. H.Y. biochemically characterized sybodies. B.Z. and Y.Z. performed neutralization assays under the supervision of D.La. N.Z., Y.G. and Q.S. performed animal experiments under supervision of Y.B. and G.W.. W.Q. collected X-ray diffraction data. B.J. helped with molecular cloning. S.K. performed half-life assays in mice. J.L. and G.W. developed reagents for the neutralizing assays. G.W. developed the mice model used for in vivo studies. D.Li. conceived the project, solved the structures, analyzed data, and wrote the manuscript with inputs from H.Y., T.L., H.C., B.Z., G.W., Y.B., M.A.S., and D.La.

## SUPPLEMENTARY MATERIALS

Materials and Methods

Extended Data Table 1-2

Extended Data Fig. 1–8

## MATERIALS AND METHODS

### Protein expression and purification – *SARS-CoV-2 S-RBD for sybody selection*

The construct for the RBD with an Avi-tag for biotinylation was made by fusing DNA, from 5’- to 3’-end, of the encoding sequence for the honey bee melittin signal peptide (KFLVNVALVFMVVYISYIYAA), a Gly-Ser linker, residues of 330-541 of the SARS-CoV-2 spike protein (Uniprot <u>P0DTC2</u>), a Gly-Thr linker, the 3C protease site (LEVLFQGP), a Gly-Ser linker, the Avi tag (GLNDIFEAQKIEWHE), a Ser-Gly linker, and a deca-His tag, into a pFastBac-backbone vector by Gibson assembly^43^. Baculoviruses were generated using standard Bac-to-Bac protocols and expression was achieved by infecting *Trichoplusia ni* High Five suspension cells at 2 × 10^6^ cells per milliliter for 48-60 h at 27 °C in flasks. The medium from 1 L of culture was filtered through a 0.22-μm membrane and incubated with 3.0 mL of Ni-Sepharose Excel (Cat 17-3712-03, GE Healthcare) in the presence of 20 mM of imidazole for 2-3 h at 4 °C with mild agitation. The beads were washed with 10 column volume (CV) of 20 mM imidazole in **Buffer A** (150 mM NaCl, 20 mM Tris HCl pH 8.0). The RBD was eluted using 300 mM of imidazole in **Buffer A**. For biotinylation^26,27^, the purified RBD with the Avi-tag intact (0.8 mg mL^−1^) was incubated with 5 mM ATP, 10 mM magnesium acetate, 43.5 μM biotin, 22 μg mL^−1^ home-purified BirA in 3.2 mL volume and incubated at 4 °C for 16 h. Biotinylated RBD was concentrated using a 10-kDa cut-off membrane concentrator to ∼3 mg mL^−1^ before loaded onto a Superdex Increase 200 10/300 GL column for size exclusion chromatography. Fractions containing the RBD were pooled, aliquoted, flash-frozen in liquid nitrogen, and stored at −80 °C before use.

### Protein expression and purification – *SARS-CoV-2 S-RBD for crystallization*

For protein crystallization, the RBD was purified as above. Both the Avi-tag and the His-tag were removed by 3C protease digestion as follows. The pooled elution from Ni-Sepharose Excel column was desalted to remove imidazole using a desalting column (Cat. 732-2010, Bio-Rad) pre-equilibrated in **Buffer A**. The desalted RBD was mixed with home-purified His-tagged 3C protease at 1:100 molar ratio (3C protease : RBD) at 4 °C for 16 h. The mixture was then passed through a Ni-NTA column which binds 3C protease, undigested RBD, and the cleaved His-tag. The flow-through fractions were collected and concentrated to 8-10 mg mL^−1^. The protein was either used directly for crystallization, or flash-frozen in liquid nitrogen and stored at −80 °C before use.

For crystallization, fresh RBD or thawed from −80 °C was mixed with desired sybodies at 1:1.5 molar ratio (RBD:sybody). After incubation on ice for 30 min, the mixture was clarified by centrifugation before size exclusion chromatography. Fractions containing the complex were pooled, concentrated to ∼10-15 mg mL^−1^ before crystallization trials.

### Protein expression and purification – *sybodies in Escherichia coli*

Sybodies were expressed with a C-terminally His-tag in *Escherichia coli* MC1061 cells. Briefly, cells carrying sybody genes in the vector pSb-init^26,27^ were grown in Terrific Broth (TB, 0.17 M KH_2_PO_4_ and 0.72 M K_2_HPO_4_, 1.2 %(w/v) tryptone, 2.4 %(w/v) yeast extract, 0.5% (v/v) glycerol) supplemented with 25 mg L^−1^ chloramphenicol to OD_600_ of 0.5 at 37 °C in a shaker-incubator at 220 rpm. The growth temperature was lowered to 22 °C and the cells were allowed to grow for another 1.5 h before induced with 0.02% (w/v) arabinose for 17 h. Cells were lysed by osmotic shock. Briefly, cells from 1 L of culture were re-suspended in 20 mL of TES-high Buffer (0.5 M sucrose, 0.5 mM EDTA, and 0.2 M Tris-HCl pH 8.0) and incubated at 4 °C for 30 min. After this dehydration step, cells were abruptly rehydrated with 40 mL of ice-cold MilliQ H_2_O at 4 °C for 1 h. The periplasmic extract released by the osmotic shock was collected by centrifugation at 20,000×g at 4 °C for 30 min. The supernatant was adjusted to contain 150 mM of NaCl, 2 mM of MgCl_2,_ and 20 mM of imidazole before added with Ni-NTA resin that had been pre-equilibrated with 20 mM of imidazole in **Buffer A** (150 mM NaCl and 20 mM Tris HCl pH 8.0). After batch-binding for 2 h, the beads were washed with 30 mM imidazole, before eluted with 300 mM imidazole in **Buffer A**. The eluted protein was either used directly or flash-frozen in liquid nitrogen and stored at −80 °C.

### Protein expression - sybody MR3-MR3-ABD in *Pichia pastoris*

The encoding gene for MR3-MR3-ABD (**Table S1**) was cloned into vector pPICZαC (Invitrogen) immediately in frame with the α-factor signal peptide. To express MR3-MR3-ABD in yeast, *Pichia pastoris* GS115 and SMD1168H were transformed with *Sac*I-linearized plasmid and selected with 0.1 and 0.5 mg mL^−1^ zeocin on an YPDS agar plate (1 %(w/v) yeast extract, 2 %(w/v) peptone, 2 %(w/v) glucose, 0.8 M sorbitol, 2 %(w/v) agarose). Colonies (12 for each strain) were inoculated into 3 mL YPD liquid medium. Cells were grown in a 30-°C incubator. After 24 h, cells were harvested, washed twice with methanol-complex medium (BMMY), and suspended in BMMY medium at a final OD_600_ of 4-5 for induction. Methanol was supplemented to the medium to 0.5 %(v/v) every 24 h. After 3 days of expression, the medium was collected by centrifugation and the secreted protein was used for SDS-PAGE analysis.

To quantify the expression level, the supernatant (10 μL) was loaded together with known amount of MR3-MR3-ABD (purified from *E. coli*, 0.5, 1, 2, 3, and 4 μg) that had been pre-mixed with medium from culture of untransformed GS115. The band intensity was semi-quantified by densitometry analysis using theImage Lab 5.2 software (Bio-Rad).

### Protein expression and purification – *divalent sybodies in mammalian cells*

The encoding sequence of MR3 was cloned into a vector harboring the hinge and Fc regions of IgG2 (**Table S1**, uniprot P01859) for secretion in mammalian cells. Expi293 cells at density of 2.3 million per milliliter were transfected with the plasmid (final concentration of 2 mg L^−1^) using linear polyethylenimine (average MW of 25 kDa, 4 mg L^−1^). Valproic acid was included at a final concentration of 2 mM. Cells were cultured in a flask for 65 h. The supernatant was collected by centrifugation and filtered through a 0.22-μm membrane. The filtrate from 2 L of culture was incubated with 3.2 mL rProtein A beads (Cat SA012005, SmartLifesciences, China) for batch binding at 4 °C for 3 h. The beads were packed into a gravity column, washed with 20 CV of PBS buffer, before eluted with 0.1 M glycine pH 3.0. The elution was quickly neutralized using 1 M Tris HCl pH 8.0. The buffer was then exchanged to PBS using a desalt column.

### Sybody selection – *ribosome display and phage display*

Sybody selection was performed using a combination of ribosome display and phage display^26,27^. *In vitro* translation of the ‘Concave’, ‘Loop’, and ‘Convex’ library was performed according to the manufacturer’s instruction (PURE*frex* 2.1 kit, Cat. PF213-0.25-EX, Genefrontier, Chiba, Japan). A reaction mix containing 1.8 μL of nuclease-free water, 4 μL of solution I, 0.5 μL of solution II, 1 μL of solution III, 0.5 μL of 10 mM cysteine, 0.5 μL of 80 mM reduced glutathione, 0.5 μL of 60 mM oxidized glutathione, and 0.5 μL of 1.875 mg mL^−1^ disulfide bond isomerase DsbC (DS supplement, Cat. PF005-0.5-EX, Genefrontier) was warmed at 37 °C. After 5 min, 0.7 μL of mRNA library, corresponding to 1.6×10^12^ mRNA molecules, was added to the pre-warmed mix for *in vitro* translation at 37 °C for 30 min. The reaction was diluted with 100 μL ice-cold **Panning Solution** (150 mM NaCl, 50 mM magnesium acetate, 0.05 %(w/v) BSA, 0.1 %(w/v) Tween 20, 0.5 %(w/v) heparin, 1 μL RNaseIn, and 50 mM Tris-acetate pH 7.4) and cleared by centrifugation at 20,000×g for 5 min at 4 °C. Biotinylated RBD was added to the supernatant and the mixture was incubated on ice for 20 min. Streptavidin beads (Dynabeads Myone Streptavidin T1) were added to pull-down the complex consisting of nascent sybody binders, the stalled ribosome with the mRNA encoding the binders, and biotinylated RBD. Selected mRNAs were purified and reverse-transcripted into single-chain DNA with the primer 5’-CTTCAGTTGCCGCTTTCTTTCTTG-3’ using a reverse transcriptase (Cat 200436, Agilent). The resulting cDNA library was purified using a DNA purification kit (Cat A740609.25, Macherey-Nagal), and PCR-amplified using the primer pair 5’-ATATGCTCTTCTAGTCAGGTTCAGCTGGTTGAGAGCG-3’ and 5’-TATAGCTCTTCATGCGCTCACAGTCACTTGGGTACC-3’ for ‘Concave’ and ‘Loop’ library, and the primer pair 5’-ATATGCTCT TCTAGTCAAGTCCAGCTGGTGGAATCG-3’ and 5’-TATAGCTCTTCATGCAGAAACGGTAACTTGGGT GCCC-3’ for the ‘Convex’ library. The product was gel-purified, digested with the Type IIS restriction enzyme *BspQ*I, and ligated into the vector pDX_init^26,27^ treated with the same enzyme. The ligation product was then transformed into *E. coli* SS320 competent cells by electroporation to generate libraries for phage display.

Three rounds of phage display were carried out. The first round was performed in a 96-well plate coated with 60 nM neutravidin (Cat. 31000, Thermo Fisher Scientific). Phage particles were incubated with 50 nM biotinylated RBD, washed, and released from the plate by tryptic digestion with 0.25 mg mL^−1^ trypsin in the buffer containing 150 mM NaCl and 20 mM Tris-HCl pH 7.4. The selected phage particles were amplified, and the second round of selection was performed by switching the immobilizing matrix to 12 μL of MyOne Streptavidin C1 beads that were pre-incubated with 50 nM biotinylated RBD. Before releasing the phage particles, the binders were challenged with 5 μM non-biotinylated RBD to compete off the binders with fast off-rates. The second selection was repeated with 5 nM of the RBD. After three rounds of selection, the phagemid was sub-cloned into pSb_init vector by fragment-exchange (FX) cloning and transformed into *E. coli* MC1061 for further screening at a single-colony level^26,27^.

### Enzyme-linked immunosorbent assay (ELISA) – *sybody selection*

Single colonies carrying sybody-encoding genes in the vector pSb-init were inoculated into 96-well plates. Cells were grown at 37 °C for 5 h in a shaking incubator at 300 rpm before 1:20 diluted into 1 mL of fresh TB medium supplemented with 25 μg mL^−1^ chloramphenicol. Cells were induced with arabinose as mentioned earlier at 22 °C for 17 h before harvested by centrifugation at 3,220 g for 30 min. Cells were resuspended in TES Buffer (20 % (w/v) sucrose, 0.5 mM EDTA, 0.5 μg/mL lysozyme, 50 mM Tis-HCl pH 8.0) and shaken for 30 min at room temperature (RT, 22-25 °C). To the lysate, 1 mL of **TBS** (150 mM NaCl, 20 mM Tris-HCl pH 7.4) with 1 mM MgCl_2_ was added. The mixtures, still in the plate, were then centrifuged at 3,220 g for 30 min at 4 °C. The supernatant containing sybodies was used directed for ELISA or FSEC assay (below).

For ELISA, Protein A was incubated with Maxi-Sorp plate 96 well (Cat. 442404, Thermo Fisher) at 4 °C for 16 h. The solution was then removed and the plate was blocked by 0.5 %(w/v) bovine serum albumin (BSA) in **TBS** buffer for 30 min at RT. The plate was washed three times using **TBS** before added with anti-myc antibodies at 1:2,000 dilution in TBS-BSA-T buffer (TBS supplemented with 0.5 %(w/v) BSA and 0.05 %(v/v) Tween 20). The antibody was allowed to bind to protein A for 20 min at RT. The plate was then washed three times with **TBST** (TBS supplemented with 0.05% Tween 20). Myc-tagged sybody prepared above was added and incubated for 20 min at RT. After washing three times with **TBST**, biotinylated RBD or MBP (the maltose-binding protein, as a control) was added to each well to a final concentration of 50 nM. After incubation for 20 min at RT, the solution was discarded and the plate was rinsed three times with **TBST**. Streptavidin conjugated with horseradish peroxidase (HRP) was added to each well (1:5,000, Cat S2438, Sigma). After incubation at RT for 30 min, the plate was washed three times again with **TBST**. ELISA signal (absorbance at 650 nm) was developed by adding 100 μL of developing buffer (51 mM Na_2_HPO_4_, 24 mM citric acid, 0.006 %(v/v) H_2_O_2_, 0.1 mg mL^−1^ 3,3’,5,5’-tetramethylbenzidine) followed by incubation at RT.

### Sybody selection – *fluorescence-detection size-exclusion chromatography (FSEC)*

To rapidly characterize RBD binders without purification, we have developed an analytic, fluorescence-detection size exclusion chromatography (FSEC)-based assay as follows. Biotinylated RBD_avi_ was bound to streptavidin (Cat 16955, AAT Bioquest) which was fluorescently labeled by fluorescein via amine coupling. The complex is named as FL-RBD_avi_. To 0.5 μM of FL-RBD_avi_, cell lysate containing unpurified sybodies were added to an estimated concentration of 0.019 mg mL^−1^, assuming expression level of 20 mg L^−1^. The mixture was loaded onto an analytic gel filtration column (Cat 9F16206, Sepax) connected to an HPLC system equipped with a fluorescence detector (RF-20A, Shimadzu). The profile was monitored by fluorescence at the excitation/emission pair of 482/508 nm. Periplasmic extract without sybodies was used as negative control. Binders can be identified based on earlier retention volume, presumably reflecting the bigger size of the FL-RBD_avi_ -sybody complex than the FL-RBD_avi_ alone.

### Bio-layer interferometry assay

The binding kinetics were measured using a bio-layer interferometry (BLI) assay with an Octet RED96 system (ForteBio). Biotinylated RBD was immobilized on a SA sensor (Cat 18-5019) that was coated with streptavidin by incubating the sensor in 2 μg mL^−1^ of RBD in **Kinetic Buffer** (0.005 %(v/v) Tween 20, 150 mM NaCl, 20 mM Tris HCl pH 8.0) at 30 °C. The sensor was equilibrated (baseline) for 120 s, before incubating with sybodies at various concentrations (association) for 120 s (for MR3) or 300 s (for all the others). The concentrations for SR4 are 0, 250, 500, 1000, and 2000 nM. The concentrations for MR17 are 0, 125, 250, 500, and 1000 nM. The concentrations for MR3/MR4 are 0, 12.5, 25, 50, and 100 nM. The sensor was then moved into sybody-free buffer for dissociation and the signal was monitored for 600 s. Data were fitted for a 1:1 stoichiometry for *K*_*D*_, *K*_on_, and *K*_off_ calculations using the built-in software Data Analysis 10.0.

For competition binding of the RBD between sybody and ACE2 (Cat 10108-H08B, Sino Biological), the RBD was immobilized and the sensor was equilibrated as abovementioned. The sensor was then saturated using 1 μM sybody and the system was equilibrated for 180 s. After saturation, the sensor was moved into sybody solutions (50 nM) with or without 25 nM ACE2. The association of ACE2 was monitored for 600 s. As a control, the ACE2-RBD interaction was monitored using sensors without sybody incubation.

For the binding assay of MR3-MR3-ABD with HSA, the sensor was coated with RBD as described earlier before saturated by incubation in 200 nM MR3-MR3-ABD before soaked with 200 nM HSA for BLI signal monitoring. A control experiment was carried out in parallel but the sensor was incubated in buffer without MR3-MR3-ABD.

### Thermostability assay

Thermostability assay of sybodies was carried out using fluorescence-detection size exclusion chromatography^44^. Sybodies at 9 μg mL^−1^ in **Buffer A** (150 mM NaCl, 20 mM Tris HCl pH 8.0) were heated at 90 and 99 °C for 20 min. The heated samples and the non-heated samples (4 °C) were analyzed the same way as described in the FSEC assay above except that the intrinsic tryptophan fluorescence (Ex. 280 nm, Em. 350 nm) was monitored.

### Pseudotyped particle production and neutralizing assays

The retroviral pseudotyped particles were generated by co-transfection of HEK293T cells using polyethylenimine with the expression vectors encoding the various viral envelope glycoproteins, the Murine leukemia virus core/packaging components (MLV Gag-Pol), and a retroviral transfer vector harboring the gene encoding the green fluorescent protein (GFP). The S Protein expressed by phCMV-SARS-CoV and phCMV-SARS-CoV-2 has been truncated in the cytoplasmic tail by adding a stop codon which removed 19 amino acids at the C-terminal. Supernatants that contained pseudotyped particles were harvested 48 h post-transfection and filtered through a 0.45-μm membrane before been used for neutralizing assays.

VeroE6-hACE2 cells (10^4^ cells/well) were seeded in a 48-well plate and infected 24 h later with 100 μL of virus supernatant in a final volume of 150 μL. Sybodies were pre-incubated with the pseudotype samples for 1 h at 37 °C prior to cell/virus co-incubation. After 6 h of co-incubation, the supernatants were removed and the cells were incubated in medium for 72 h at 37 °C. GFP expression was determined by fluorescence-activated flow cytometry analysis. The infectivity of pseudotyped particles incubated with sybodies was compared with the infectivity observed using pseudotyped particles and Dulbecco’s modified Eagle’s medium-2% fetal calf serum only and standardized to 100%.

Average and standard deviation (SD, n=3) were plotted for the IC_50_ experiments except for Fig. 4D which reports data from two independent experiments.

### Crystallization

Crystallization trials were set up using a Crystal Gryphon LCP robot as follows. To a two-well sitting-drop plate, 70 μL of precipitant solution was added to the reservoir. To each well, 150 nL of protein solution was added using the LCP arm of the robot. The wells were covered with 150 nL of precipitant solution using the 96-headed tips. Plates were sealed using a tape (Cat HR4-506, Hampton research) and placed at 20 °C in a Rocker Imager 1000 for automatic imaging.

Crystals for the SR4-RBD complex were grown in 20% (w/v) PEG 3,000, 200 mM sodium chloride, 100 mM HEPES pH 7.5. Cryo protection was achieved by adding 20 %(v/v) glycerol to the mother liquor condition. Crystals for the MR17-RBD complex were grown in 20 %(w/v) PEG 3,350, 0.2 M magnesium formate. Cryo protection was achieved by adding 10 %(v/v) glycerol in the mother liquor condition. Crystals for the MR3-RBD complex were obtained in 9 %(w/v) PEG 8,000, 0.1 M HEPES pH 7.5, 8 %(v/v) ethylene glycol, 9.6 %(v/v) glycerol. 20% glycerol was included for cryo cooling. Crystals for the MR4-RBD complex were grown in 10 %(w/v) PEG 8,000, 200 mM zinc acetate, 100 mM MES pH 6.0. Crystals for MR17-K99Y were grown in 0.2 M MgCl_2_, 20 %(w/v) PEG 3,350. Cryo protection was performed by adding 30 %(v/v) glycerol to the reservoir condition. Crystals were cryo-protected, harvested using a MitGen loop, and flash-cooled in liquid nitrogen before X-ray diffraction data collection.

### Data collection and structure determination

X-ray diffraction data were collected at beamline BL19U1 (ref.^45^) at Shanghai Synchrotron Radiation Facility. Diffraction data were collected with a 50 x 50 μm beam on a Pilatus detector at a distance of 300 – 500 mm, with oscillation of 0.5 - 1° and a wavelength of 0.97853 Å. Data were integrated using XDS ^46^, and scaled and merged using Aimless ^47^. The structure was solved by molecular replacement using Phaser ^48^ with the RBD structure from PDB 6M0J and the sybody from PDB 5M13 ^26^ as the search model. The model was built with 2F_o_-F_c_ maps in Coot ^49^, and refined using Phenix ^50^. Structures were visualized using PyMol ^51^.

### Structure-based design of sybody mutants to improve binding affinity

The structure of the MR17-RBD complex was examined using Coot^49^ and PyMol^51^. A panel of 19 single mutants was designed by virtual mutation using Coot ^49^ followed by examining the possible increasing in numbers of H-bonds, salt bridges, or hydrophobic interactions. The mutations include V31F, V31I, E35F, G47A, G47F, G47W, E52F, E52M, E52Q, S53k, S53Q, H56F, H56I, H56W, H56Y, K99Y, Q103D, Q103E, and Q103Y. The mutants were purified and characterized the same way as for MR17. Because K99Y showed higher neutralization activity than the wild-type, K99W was designed for the second round.

### *In vivo* stability of sybody in mice

The female 7-week-old ICR mice weighing 27 ± 1 g were intraperitoneally injected with phosphate buffered saline (PBS) or sybodies MR3, MR3-MR3, or MR3-MR3-ABD at 25mg kg^−1^ in a final volume of 100 μL in PBS. The blood samples were collected at different time points (2 days preinjection, 6 h, 12h, 1 day, 3 days, 6 days and 14 days postinjection) and subjected to neutralization assay using SARS-CoV-2 pseudotypes. Mice weights were measured till 6 days post-injection (n=4). Mice were sacrificed at 1, 3, and 6 days post-injection; their vital organs (heart, liver, spleen, lung, kidney and thymus) were fixed in 4% formaldehyde at 4 °C overnight and then embedded within paraffin, solidified and cut to 15-μm thickness using a cryotome (Leica Microsystems). Sections were stained by hematoxylin and eosin. Scale is equal to the original magnification ×100.

### Mice challenge experiments

C57BL/6J female mice (6-8 weeks old) were treated with adenovirus serotype 5 expressing human angiotensin 1 converting enzyme 2 (hACE2) via the intranasal route as previously described^40^. At 5 days post-adenovirus treatment, the mice were intranasally infected with SARS-CoV-2 strain hCoV-19/China/CAS-B001/2020 (National Microbiology Data Center *NMDCN0000102-3*, GISAID databases *EPI_ISL_514256-7*) with a high dose of 5 × 10^6^ TCID_50_ in a volume of 50 μL. After 12 h, the mice of MR3-MR3-ABD group (n=6) was given 200 μL of sybody each (25 mg kg^−1^ body weight) by intraperitoneal injection. The infection control group (n=3) was treated with PBS buffer. Three days post-infection (d.p.i), three mice were euthanized, and the lung tissues (∼1/8 of the total lungs) were fixed in 4 %(v/v) paraformaldehyde for histopathological analysis using hematoxylin-eosin staining. The rest of the lungs were weighted and homogenized for RNA extraction and virus titration by quantitative reverse transcription PCR (qRT-PCR) using a kit (Mabsky Biotech Co., Ltd.) following manufacturer protocols. Average and standard deviation of all three individual data points were reported.

### Ethics Statement

The animal experiments were approved by the Institutional Animal Care and Use Committee of the Institut Pasteur of Shanghai, Chinese Academy of Sciences (Animal protocol No. A2020009) for in vivo stability assays, and by the Ethics Committees of Institute of Microbiology, Chinese Academy of Sciences (SQIMCAS2020010) for the live virus-related work. The study was conducted in strict accordance with the recommendations provided in the Guide for the Care and Use of Laboratory Animals of the Ministry of Science and Technology of the People’s Republic of China. All experiments with live viruses and animals were performed in a biosafety level 3 laboratory and complied with the instructions of the institutional biosafety manual.

## Data availability

The structure factors and coordinates are available through the protein data bank (PDB) under accession codes 7C8V (SR4-RBD), 7C8W (MR17-RBD), and 7CAN (MR17-K99Y in complex with the RBD).

^*a*^Fluorescence-detection size exclusion chromatography (FSEC) assay for RBD binders. Periplasmic extraction was directly mixed with 0.5 μM of fluorescently labeled RBD and the mixture was loaded onto an analytic gel filtration column. Sybodies that caused earlier retention volume (peak shift) are labeled ‘Y’ and colored red. Sybodies that did not peak shift are indicated with ‘N’. Sybodies that were not determined for FSEC peak-shift are labeled with ‘N.D.’. ^*b*^The sequences include ‘GSSS’ at the N-terminal, and ‘AGRAG*EQKLISEEDL*NSAVDHHHHHH’ at the C-terminal which contains a myc-tag (italic) for ELISA and a hexahistidine tag for purification. GS linker are highlighted in italic, when applicable.

**Table S1.**
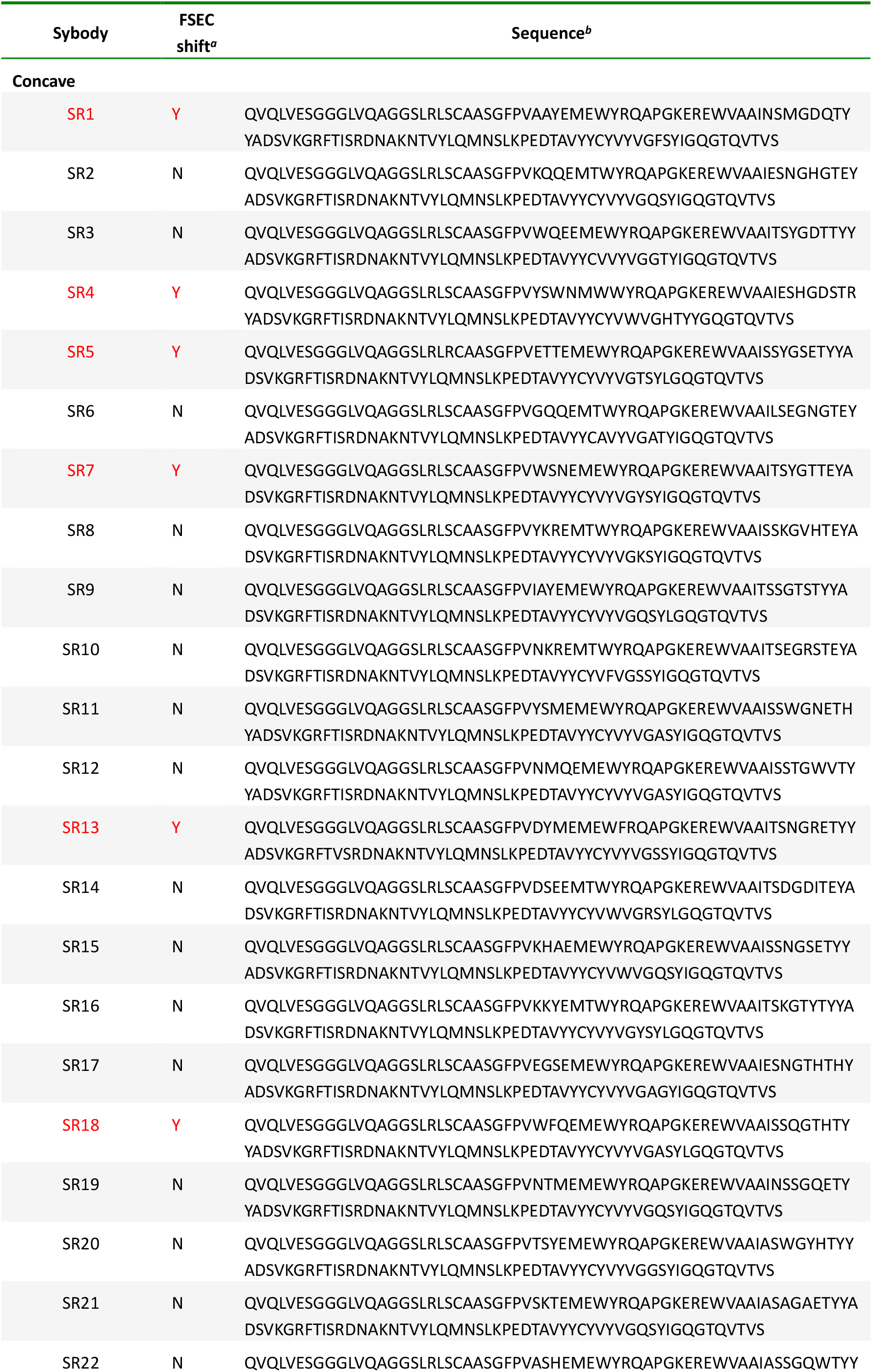
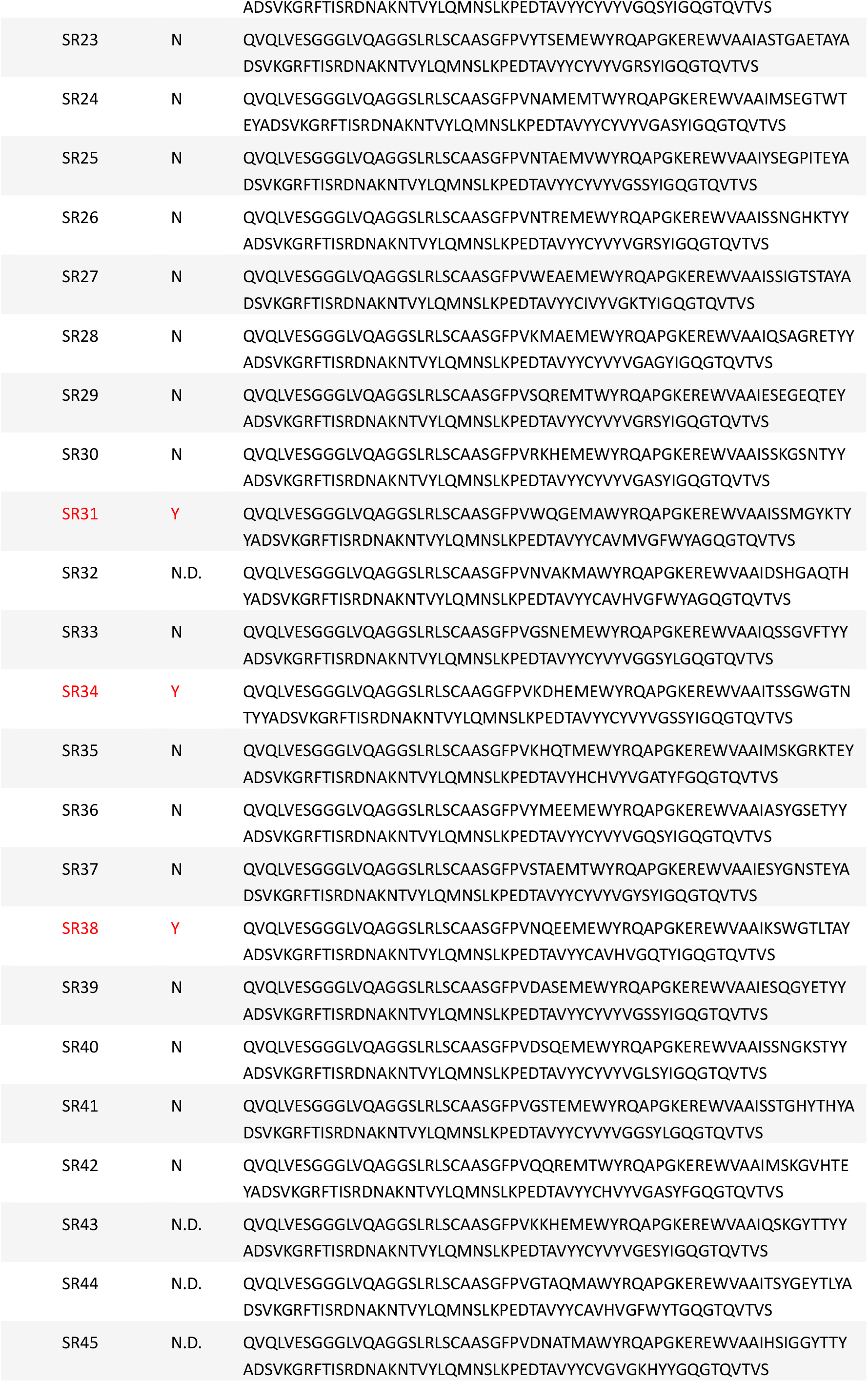
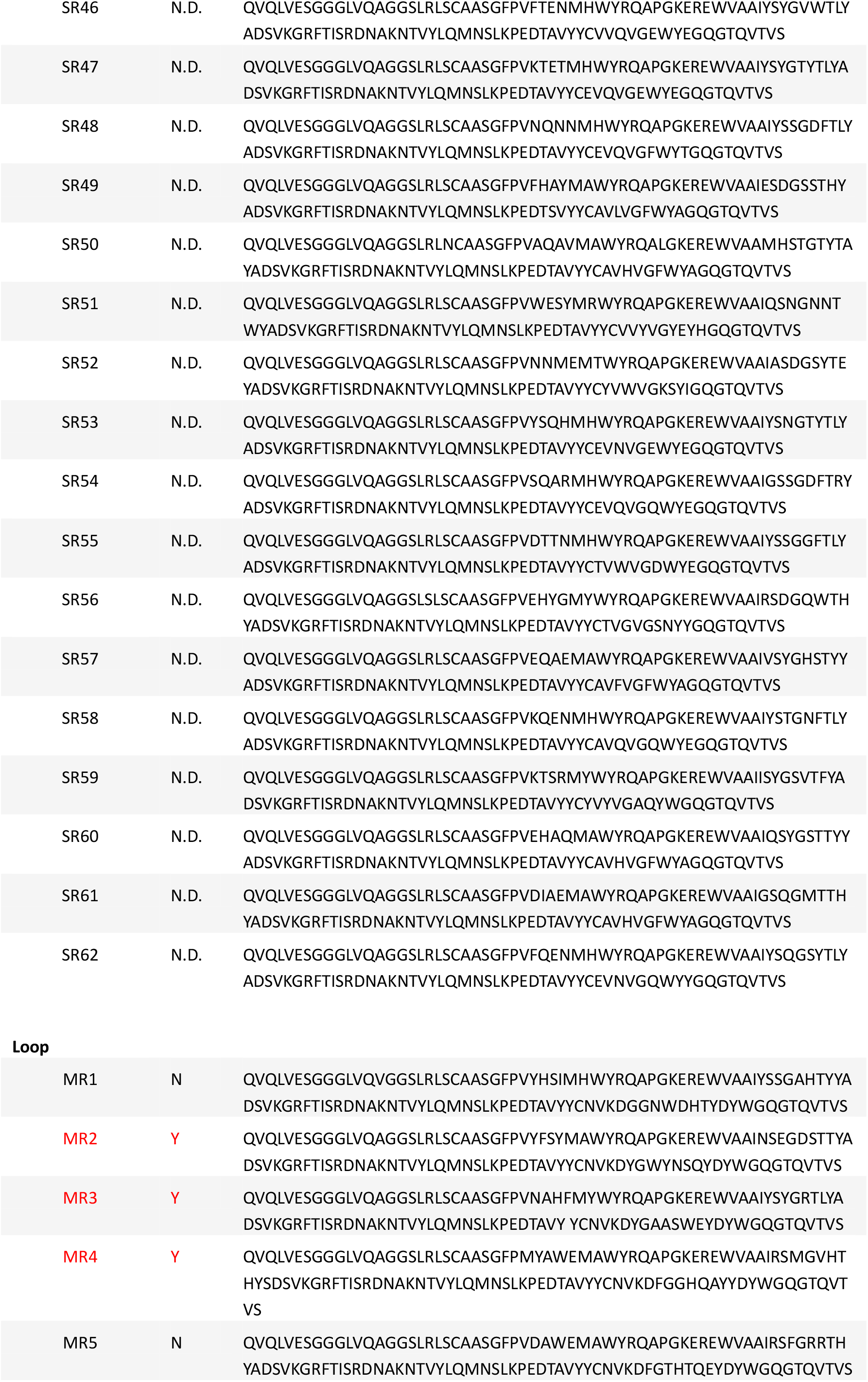
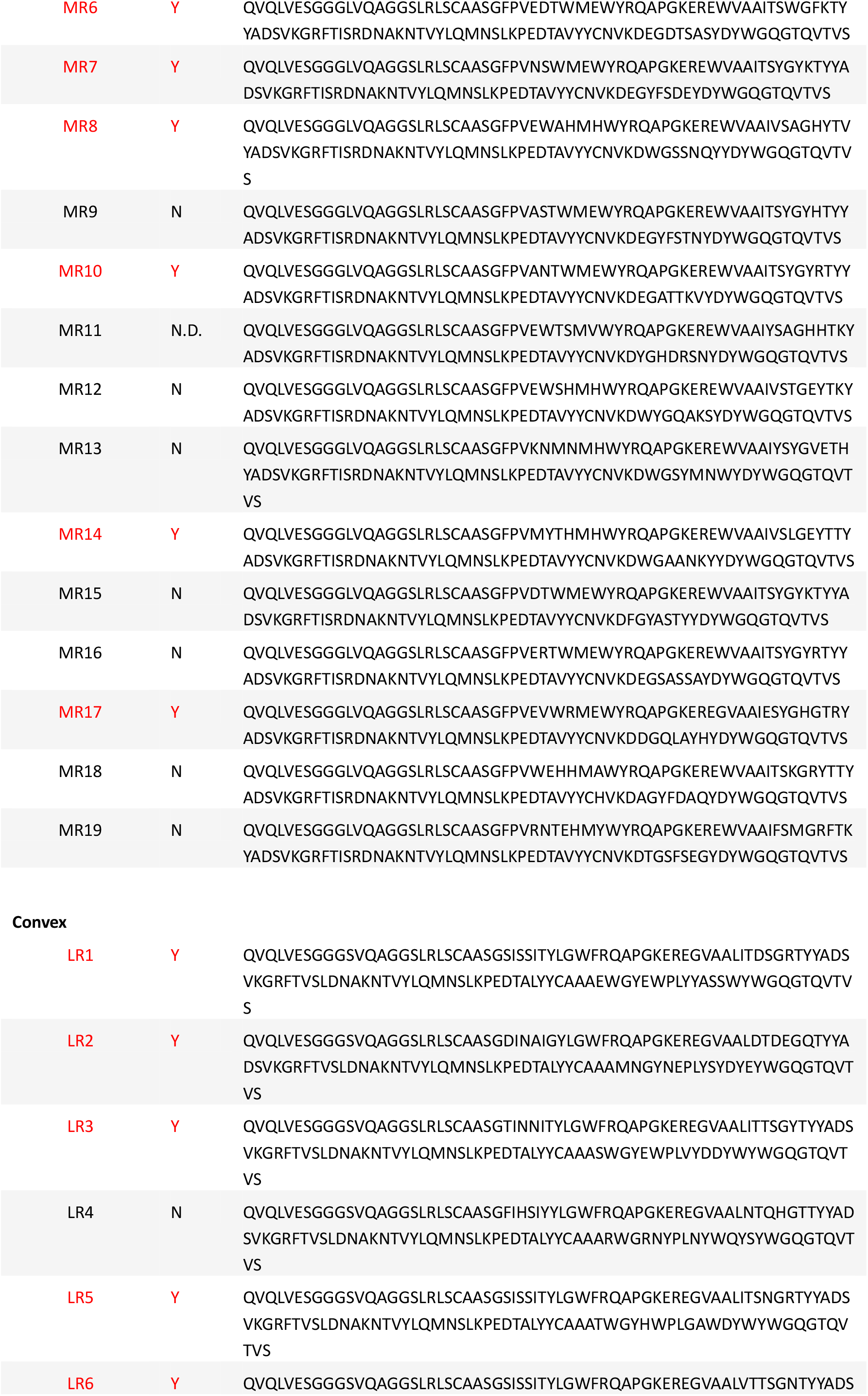
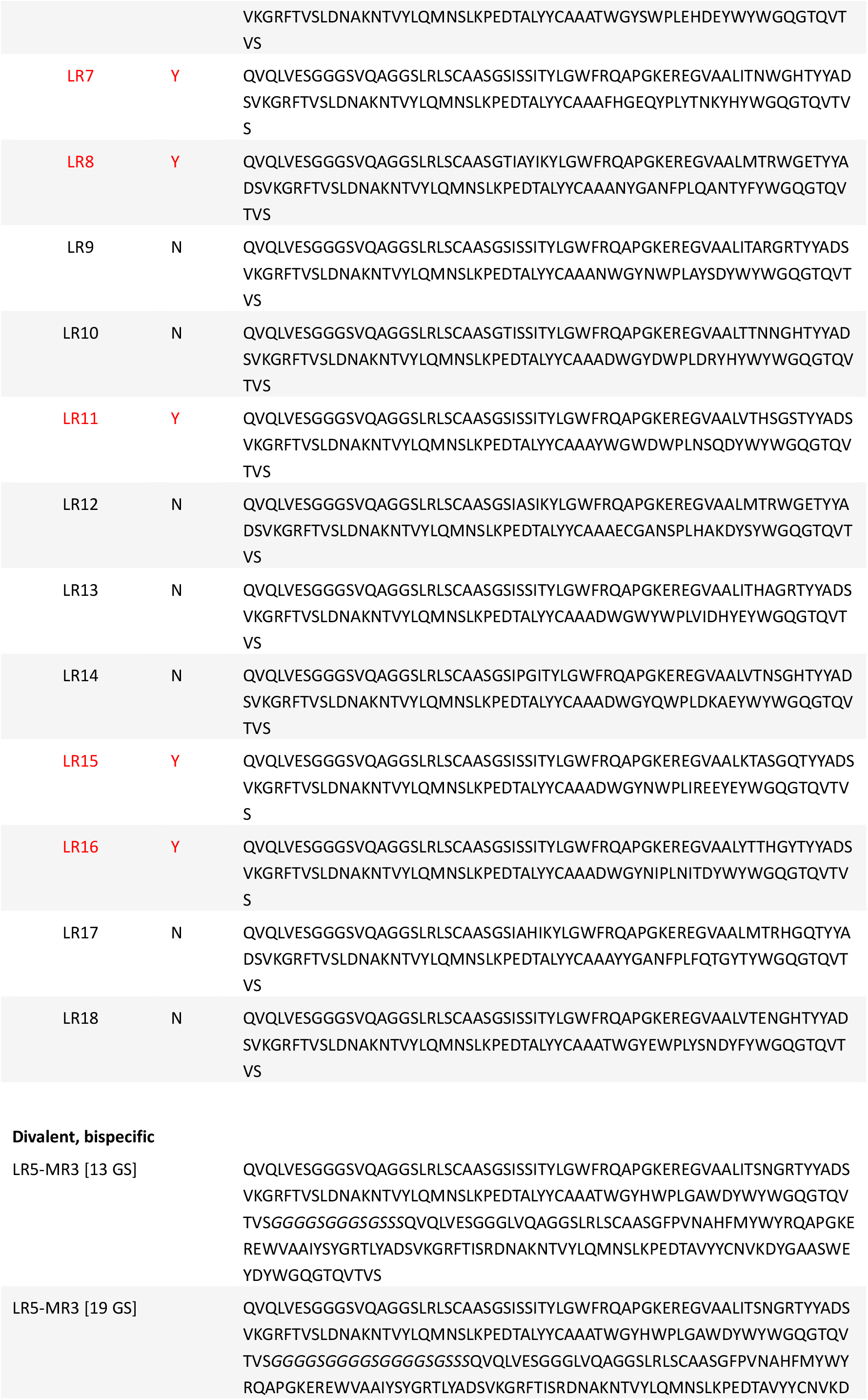
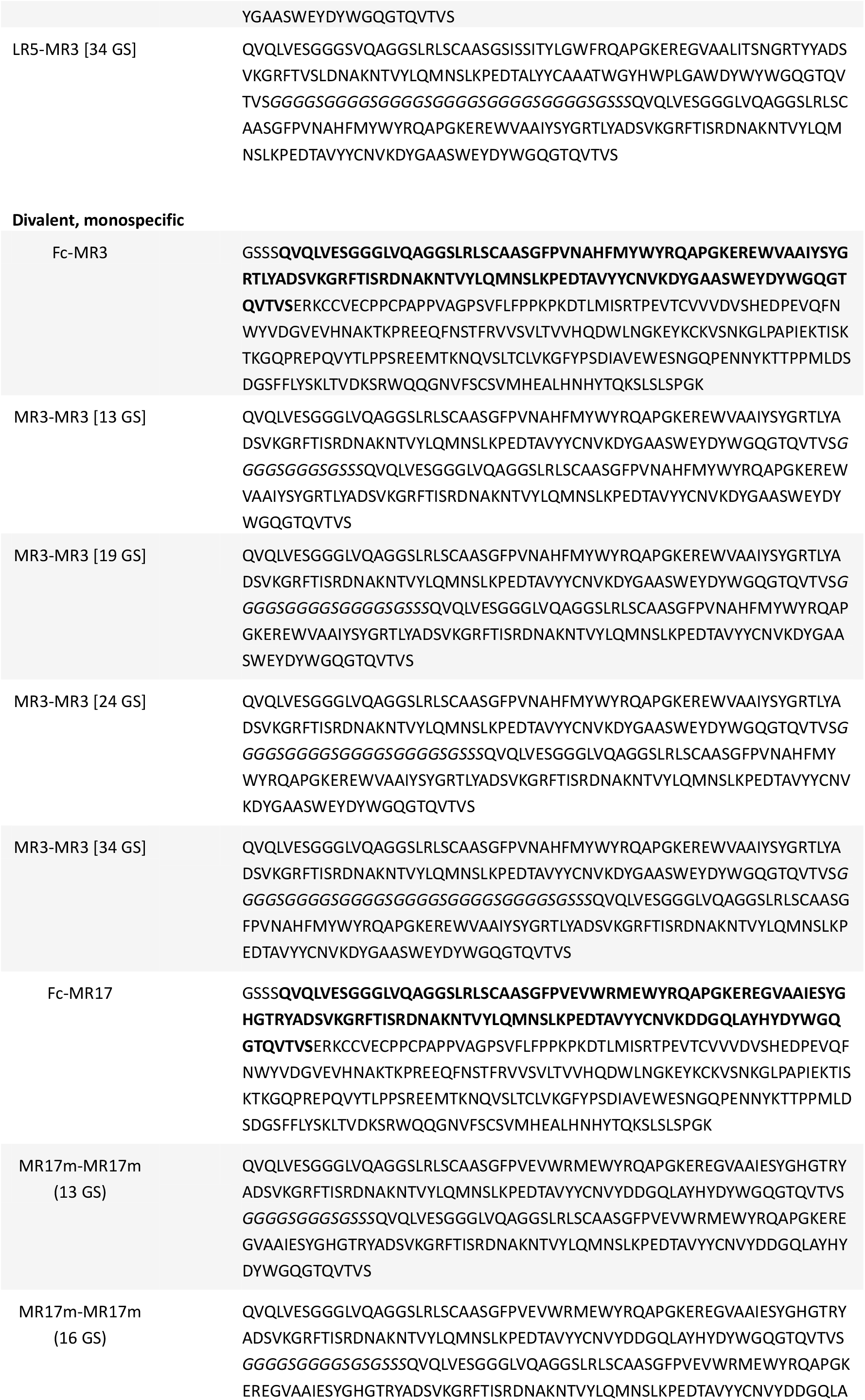
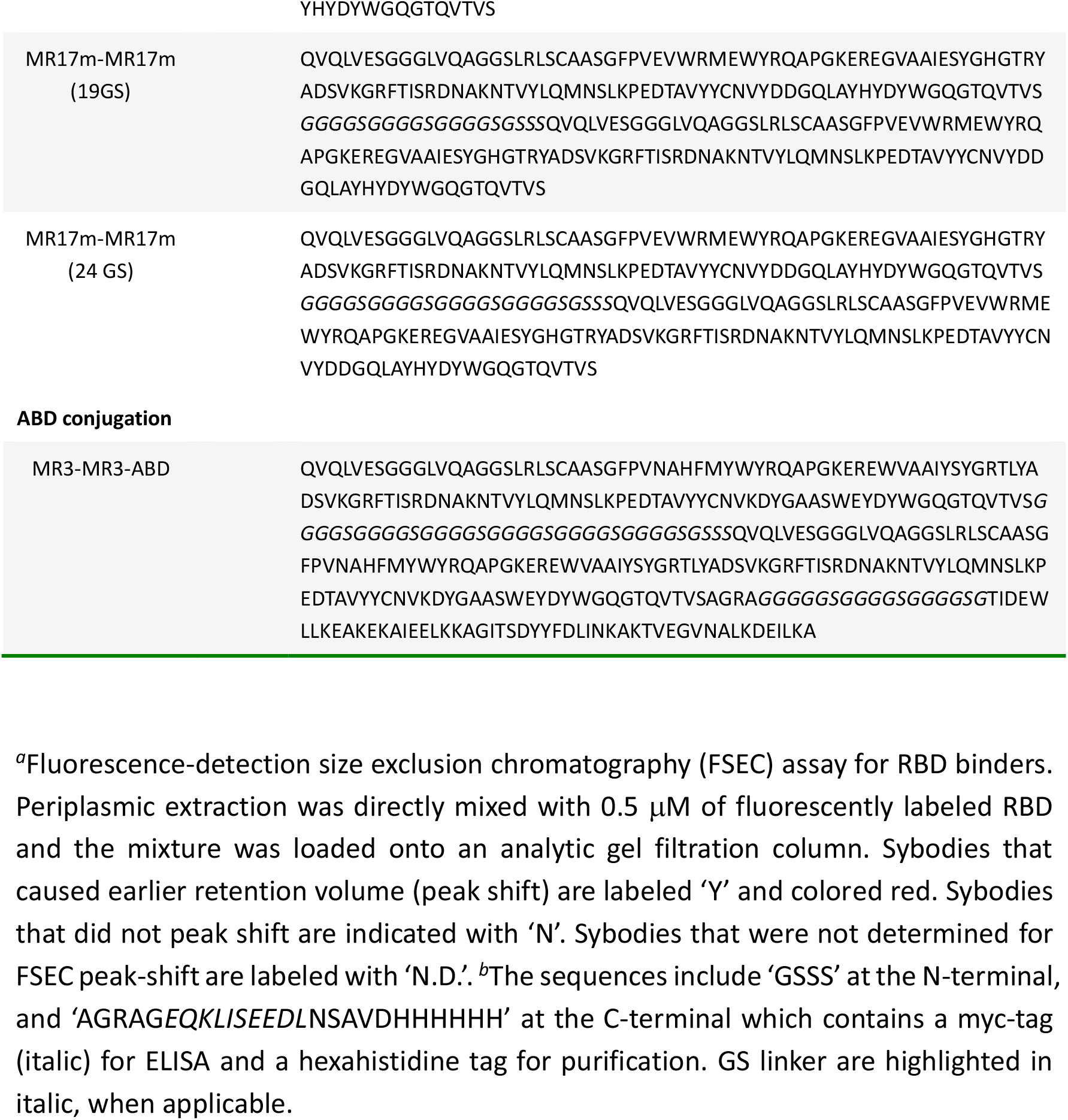
Sequences and FSEC results of sybody binders for the SARS-CoV-2 RBD.

**Table S2.** Data collection and refinement statistics.

|  | SR4-RBD | MR17-RBD | MR17(K99Y)-RBD |
| --- | --- | --- | --- |
| **Data collection** |  |  |  |
| Space group | P 6<sub>5</sub> 2 2 | P3<sub>2</sub> 2 1 | P3<sub>2</sub> 2 1 |
| Cell dimensions |  |  |  |
| *a*, *b*, *c* (Å) | 65.55, 65.55, 344.53 | 73.69, 73.69, 158.58 | 74.19, 74.19, 158.40 |
| *α*, *β*, *γ* (°) | 90, 90, 120 | 90, 90, 120 | 90, 90, 120 |
| Wavelength (Å) | 0.97853 | 0.97853 | 0.97853 |
| Resolution (Å) | 47.40 - 2.15<br>(2.23 - 2.15)<sup>a</sup> | 49.71 - 2.77<br>(2.89 - 2.77) | 49.90 - 2.94<br>(3.12 - 2.94) |
| *R*<sub>merge</sub> | 0.161 (1.203) | 0.276 (2.222) | 0.218 (1.666) |
| *R*<sub>pim</sub> | 0.054 (0.395) | 0.062 (0.494) | 0.052 (0.385) |
| *I*/*σI* | 11.4 (2.0) | 11.1 (1.5) | 12.4 (1.9) |
| Completeness (%) | 99.8 (99.9) | 100 (99.9) | 99.9 (99.6) |
| Multiplicity | 9.5 (9.9) | 21.0 (20.8) | 18.4 (19.3) |
| *CC* * <sup>b</sup> | 0.999 (0.970) | 0.997 (0.927) | 0.998 (0.920) |
| **Refinement** |  |  |  |
| Resolution (Å) | 47.40 - 2.15 | 49.71 - 2.77 | 49.90 - 2.94 |
| No. reflections | 25,148 | 13,256 | 11,264 |
| *R*<sub>work</sub> / *R*<sub>free</sub> | 0.1836 / 0.2239 | 0.2029 / 0.2659 | 0.2149 / 0.2676 |
| No. atoms | 2,810 | 2,536 | 2,509 |
| Protein | 2,510 | 2,482 | 2,465 |
| Ligands | 62 | 54 | 44 |
| Water | 238 | 0 | 0 |
| No. residues | 322 | 315 | 312 |
| B-factors (Å<sup>2</sup>) | 35.13 | 73.28 | 79.38 |
| Protein | 34.16 | 72.27 | 78.53 |
| Ligand/ion | 55.11 | 119.79 | 126.90 |
| Water | 40.22 |  |  |
| R.m.s deviations |  |  |  |
| Bond lengths (Å) | 0.007 | 0.011 | 0.004 |
| Bond angles (°) | 0.850 | 1.10 | 0.68 |
| Ramachandran |  |  |  |
| Favoured (%) | 98.06 | 96.12 | 96.08 |
| Allowed (%) | 1.94 | 3.88 | 3.59 |
| Outlier (%) | 0 | 0 | 0.33 |
| **PDB ID** | 7C8V | 7C8W | 7CAN |

<sup>a</sup>Highest resolution shell is shown in parenthesis. <sup>b</sup> $CC^* = \sqrt{\frac{2CC_{1/2}}{1+CC_{1/2}}}$

**Extended Data Fig. 1.**
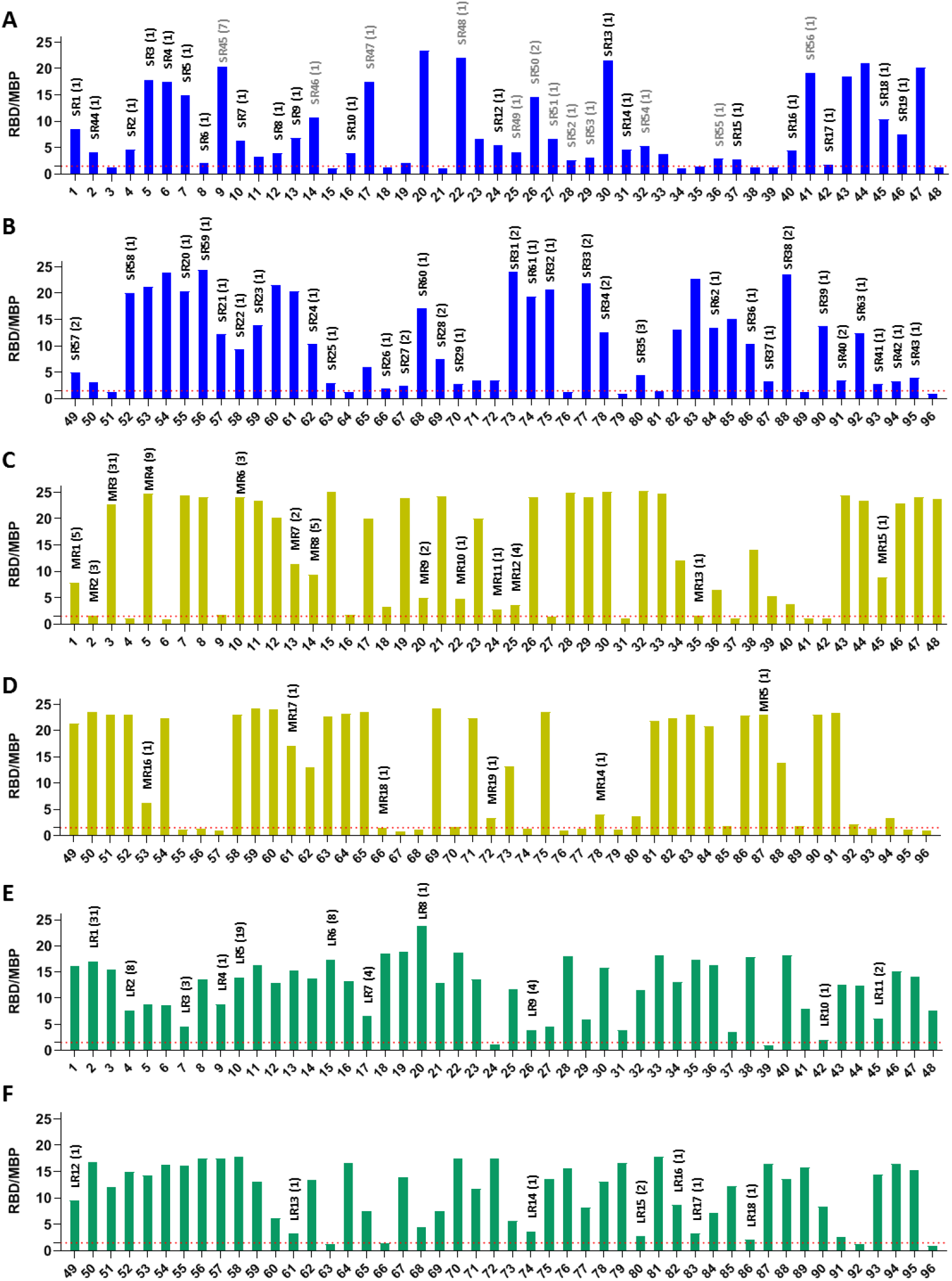
Identification of RBD binders using ELISA. (**A**,**B**) Results for the Concave library. (**C**,**D**) Results for the Loop library. (**E**,**F**) Results for the Convex library. The ratio between the ELISA signal (A_650_) of wells with the RBD and of wells with the unrelated maltose-binding protein (MBP) is plotted. The signal for MBP is typically between 0.04-0.09. A red dashed line guides the cut-off at a ratio of 1.5. Unique clones are labeled with the redundancy shown in brackets.

**Extended Data Fig. 2.**
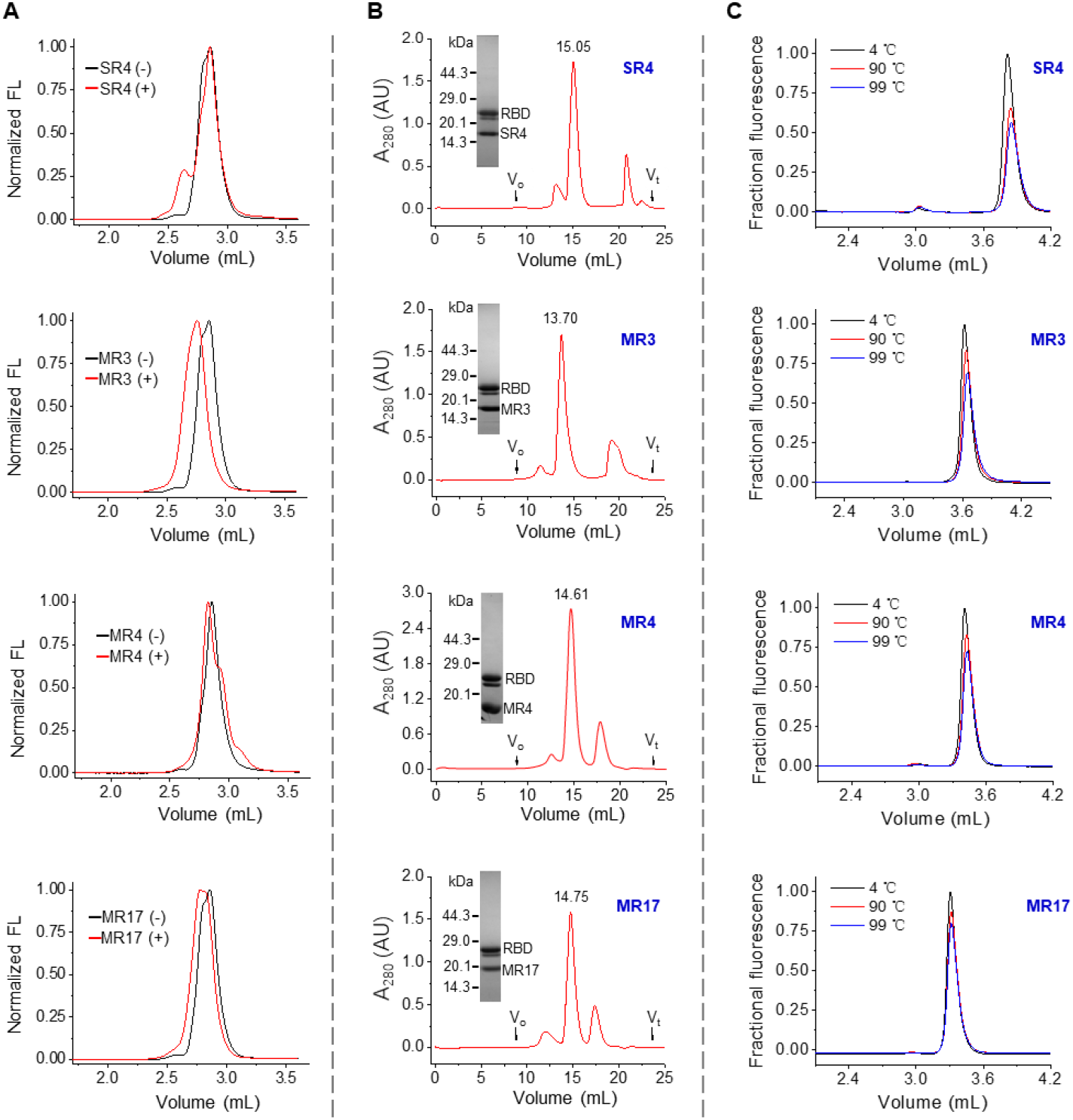
Characterization and purification of sybody-RBD complexes. (**A**) Fluorescence-detector size exclusion chromatography (FSEC) of the RBD in the absence (black, −) and presence (red, +) of crude periplasmic extract from sybody clones. Biotinylated RBD was fluorescently labeled through binding to streptavidin that was conjugated with an amine-reactive fluorescein variant. The concentration of RBD was 0.5 μM. Fluorescence (Ex. 482 nm, Em. 508 nm) was normalized before plotting. The extent of peak shift follows the order of SR4<MR4<MR17<MR3. Fluorescence trace before the void volume (V_o_, 1.78 mL) is not shown. (**B**) Preparative size exclusion chromatography of the indicated sybody-RBD complexes. SDS-PAGE images of the main-peak fraction for all four sybodies are shown in the inset. Numbers label the elution volume for the main peak. The results for SR34, SR38, MR6, LR1, and LR5 were similar to the 4 sybodies here and are not shown. (**C**) Fluorescence-detection size exclusion chromatography (FSEC) profile of the thermostability assay. Sybodies SR4, MR17, MR3, and MR4 were incubated at indicated temperatures for 20 min before loading on to an analytical size exclusion chromatography column. The elution profile was monitored using the intrinsic tryptophan fluorescence. Fluorescence intensities were normalized to the peak value of the unheated sample.

**Extended Data Fig. 3.**
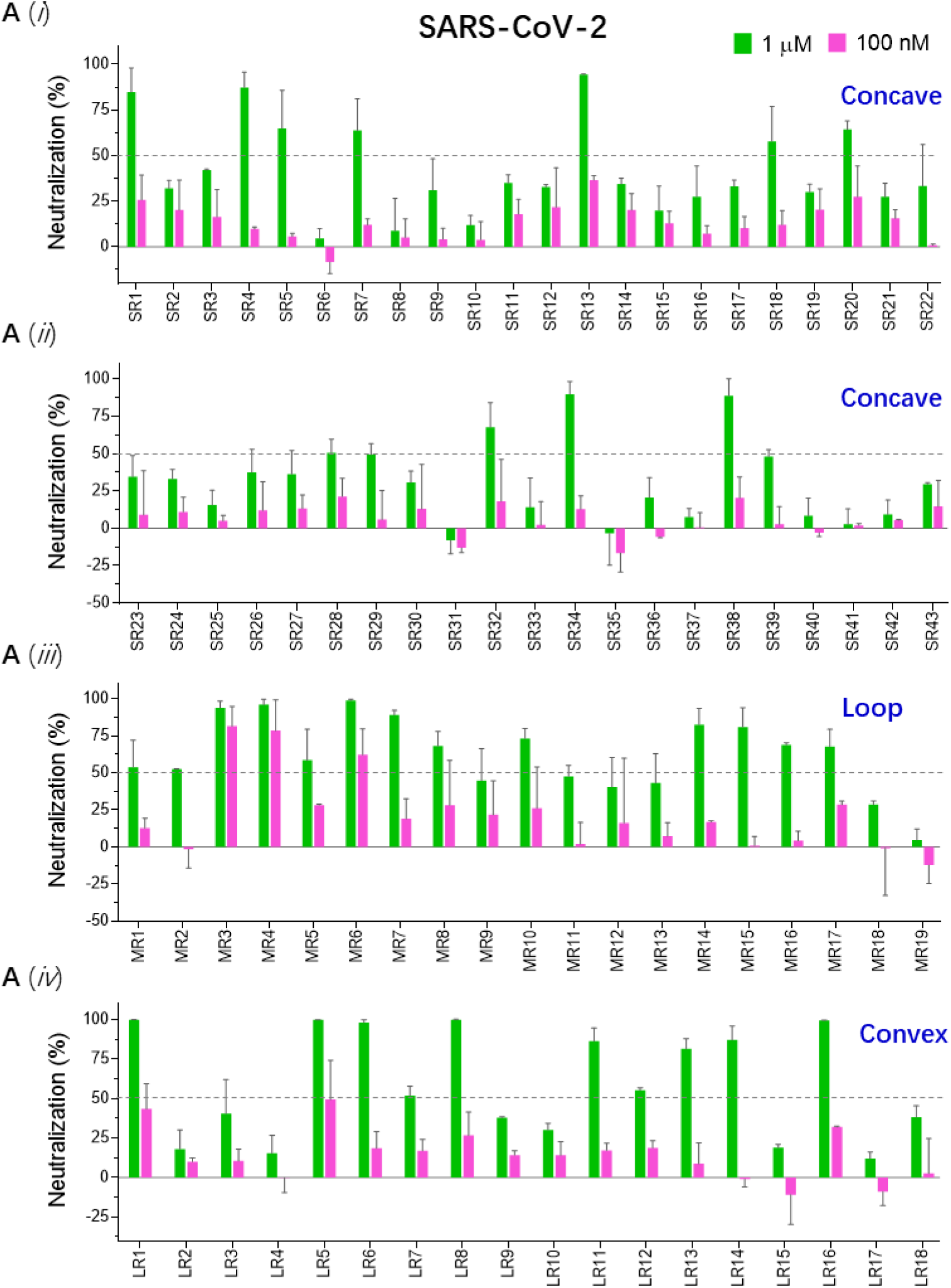
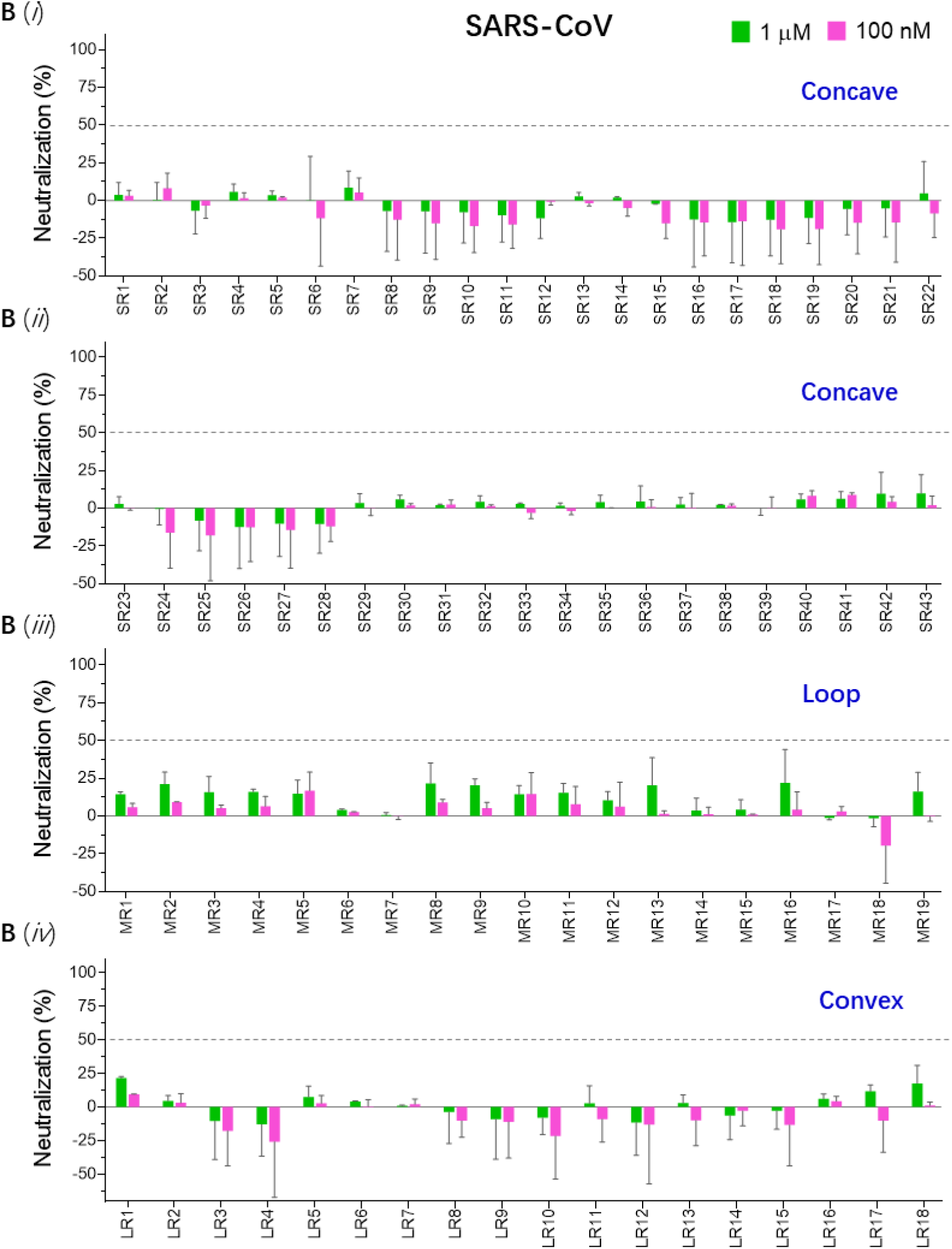
Neutralization activity of 80 sybodies. (**A**) Neutralization assay results for SARS-CoV-2 pseudovirus. (**B**) Neutralization assay results for SARS-CoV pseudovirus. VeroE6-hACE2 cells were infected with a premix of pseudotypes and sybodies at two concentrations (1 μM and 100 nM). Infectivity were measured after 3 days using FACS and the percentage of neutralization was calculated for each sybody.

**Extended Data Fig. 4.**
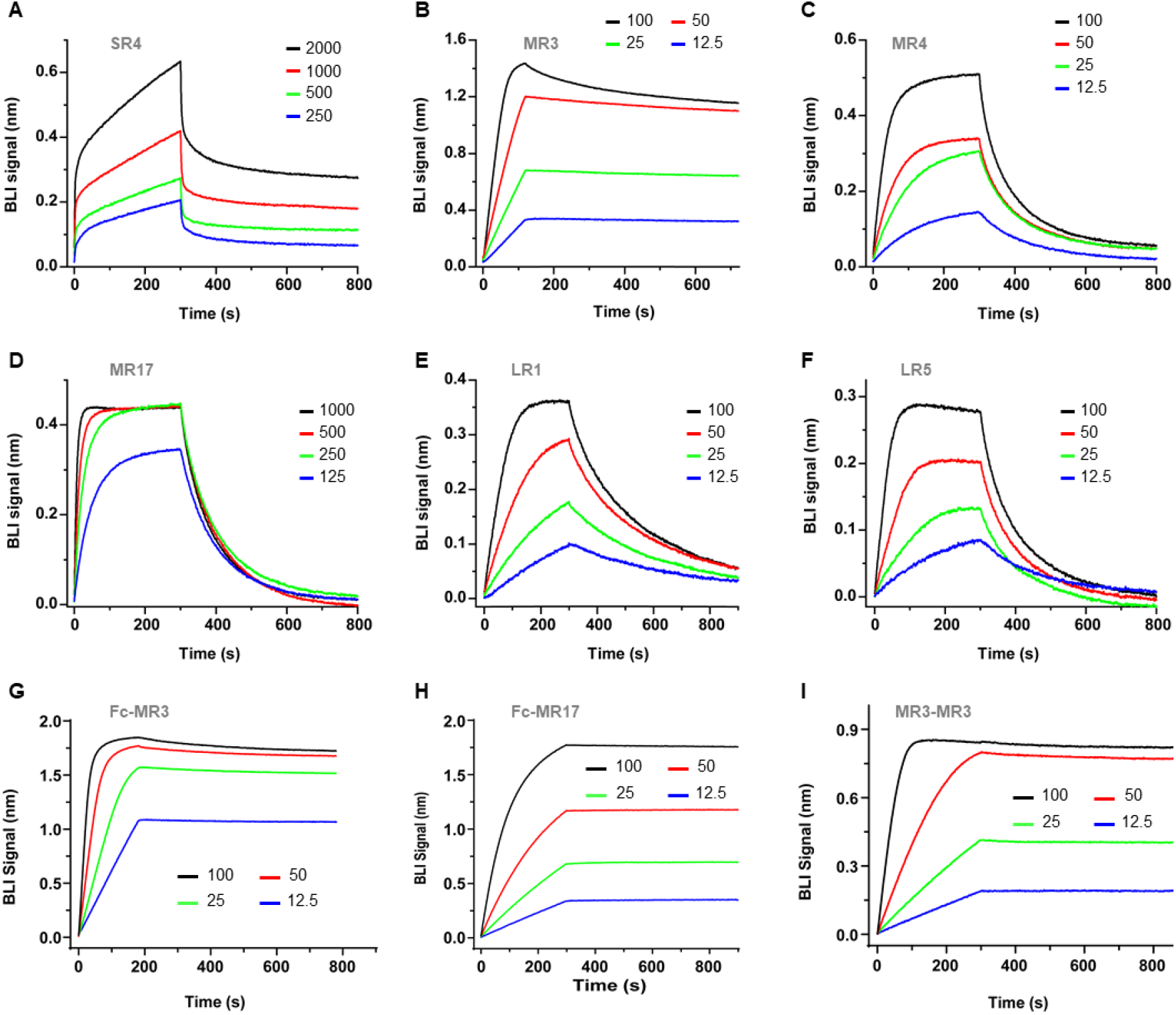
Kinetics for sybody-RBD binding. (**A-I**) Biotinylated RBD immobilized on a streptavidin-coated sensor was titrated with various concentrations (nM) of sybodies as indicated. Bio-layer interferometry (BLI) data were fitted with a 1:1 stoichiometry.

**Extended Data Fig. 5.**
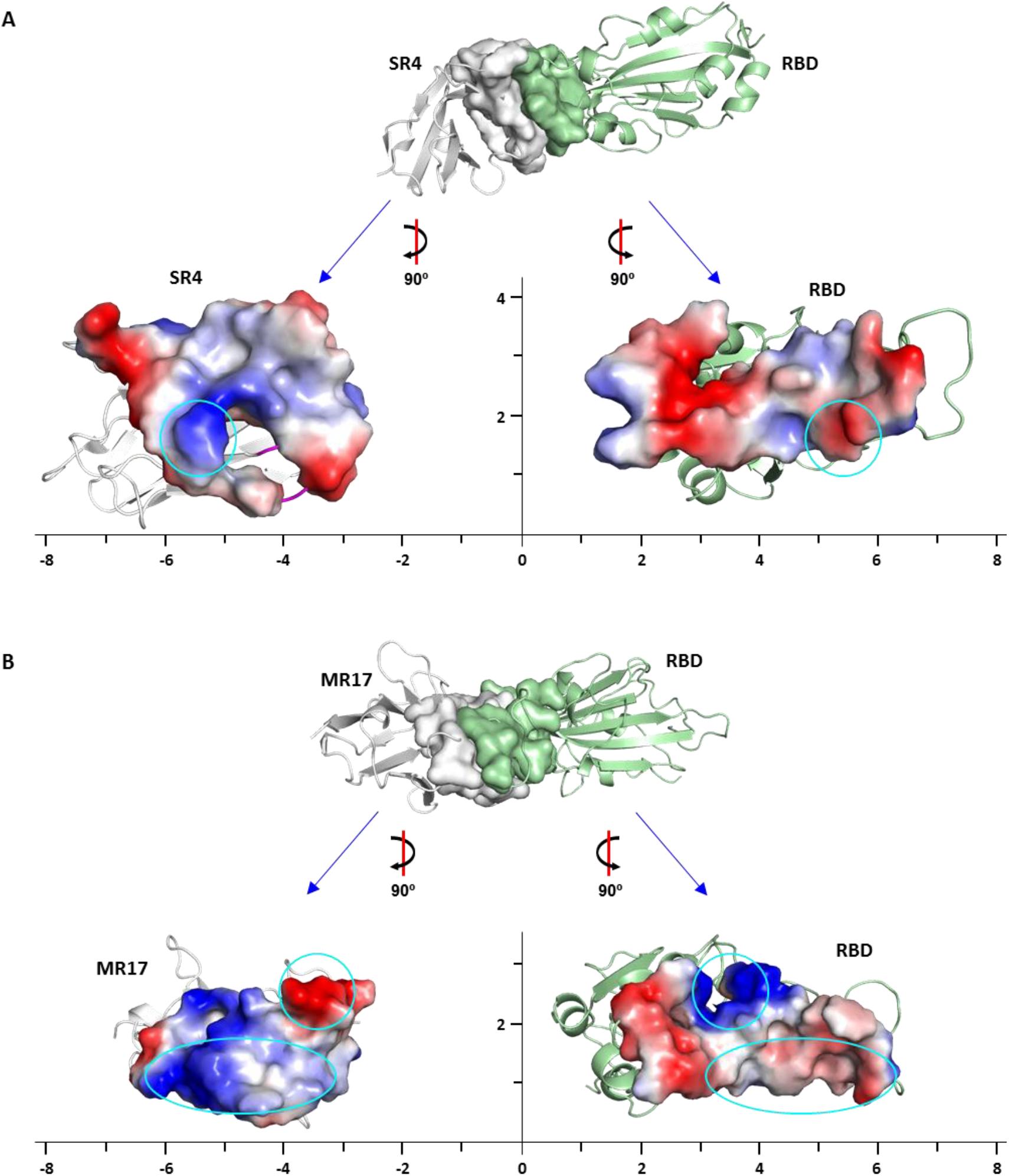
Electrostatic complementarity of the sybody-RBD binding surface. (**A**,**B**) ‘Open-book’ view of molecular electrical potential surfaces of the interface between the RBD and SR4 (**A**) and between the RBD and MR17 (**B**). The electrical potential maps were calculated by Adaptive Poisson-Boltzmann Solver (APBS) ^52^ built-in in PyMol. The unitless ruler guides the view of the relative distances between the opened surface pairs. Cyan circles highlight electrostatic complementarity.

**Extended Data Fig. 6.**
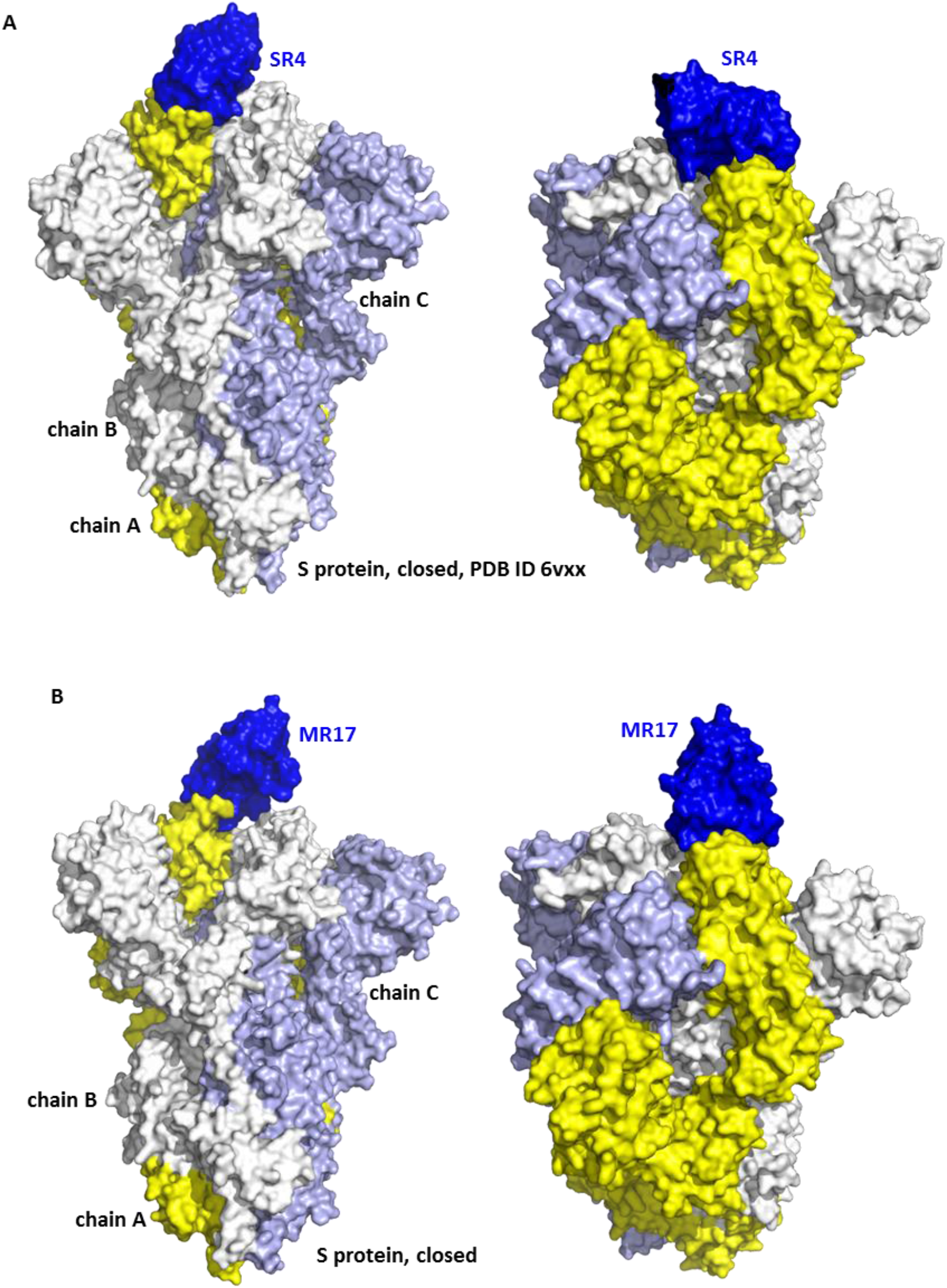
SR4 and MR17 may bind to the SARS-CoV-2 S RBD in the ‘closed’ conformation. (**A**,**B**) The structure of SR4-RBD (**A**) and MR17-RBD (**B**) were aligned to the closed conformation (PDB ID 6VXX)^2^ of SARS-CoV-2 S protein. No significant clashes were observed for both sybodies. The three chains of S are colored yellow, white, and pale blue. Sybodies are colored blue.

**Extended Data Fig. 7.**
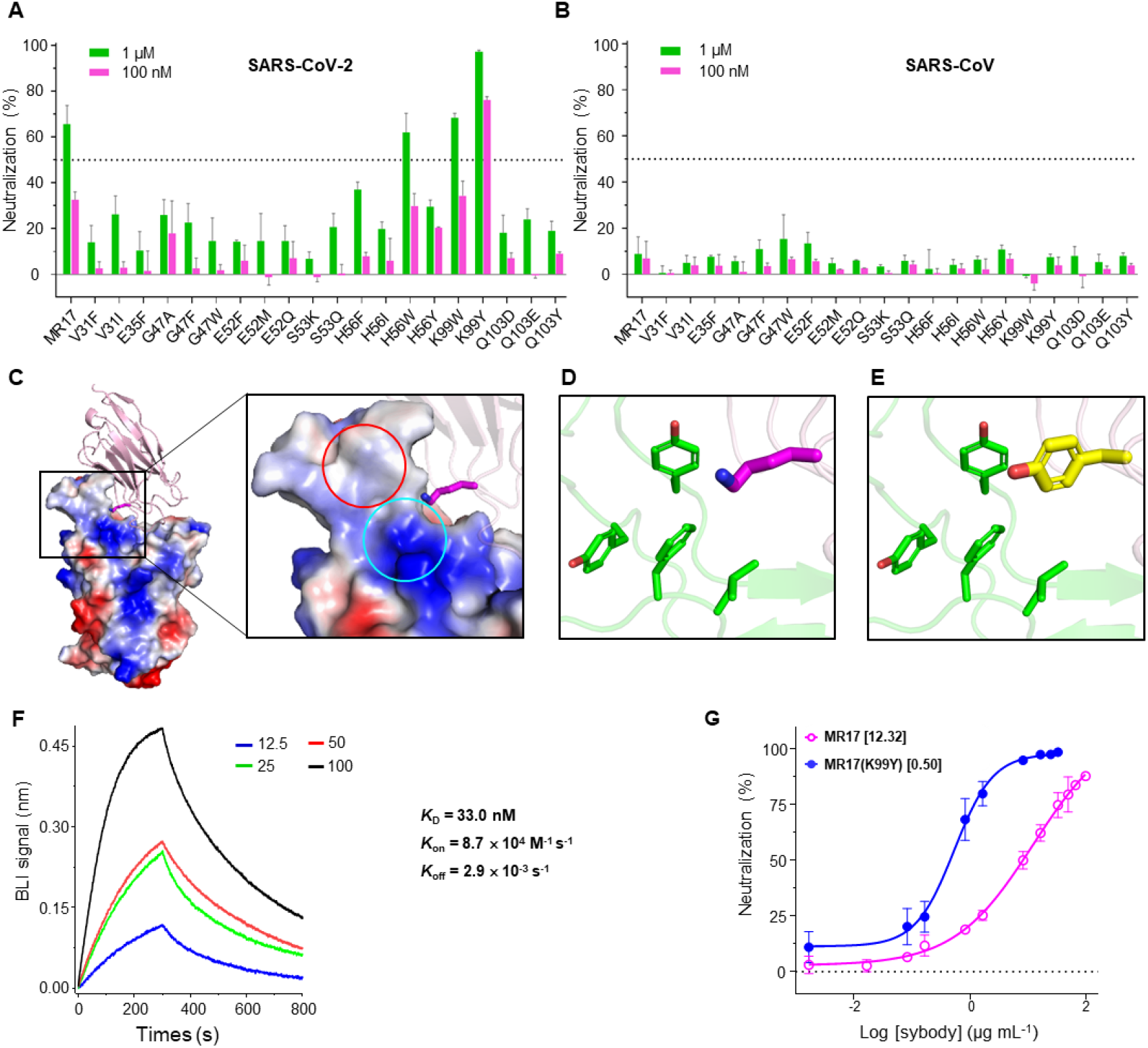
Structure-based design of a MR17 mutant (MR17m) with improved affinity and potency. (**A**,**B**) Neutralization assay for SARS-CoV-2 (**A**) or SARS-CoV (**B**) pseudotypes by the wild-type MR17 sybody and the 20 rationally designed single-mutants (See **Methods**). Sybody concentrations were used at 1 μM (green) and 100 nM (magenta) concentrations. Data are from three independent experiments. (**C**,**D**) Rational for the design of K99Y. The positively charged Lys99 pokes to an area (boxed) that contains a hydrophobic patch (red cycle) and a positively-charged surface (cyan cycle). Electrostatic repel and hydrophobic mismatch would make Lys99 unfavorable at this position. According to the original library design, Lys99 was unvaried^26^, meaning that Lys99 was not *selected* and hence opportunities for optimization. (**E**) The K99Y mutation fits the hydrophobic microenvironment well, as revealed by the crystal structure of MR17m (**Extended Data Table 2**). (**F**) Binding kinetics of MR17m binding to RBD. BLI signals were recorded under indicated MR17m concentrations (nM). (**G**) Comparison of neutralization activity of MR17 and MR17m. IC_50_ values (μg mL^−1^) for SARS-Cov-2 are indicated in brackets. Data for MR17 are from **Fig. 1B**. Data are from three independent experiments.

**Extended Data Fig. 8.**
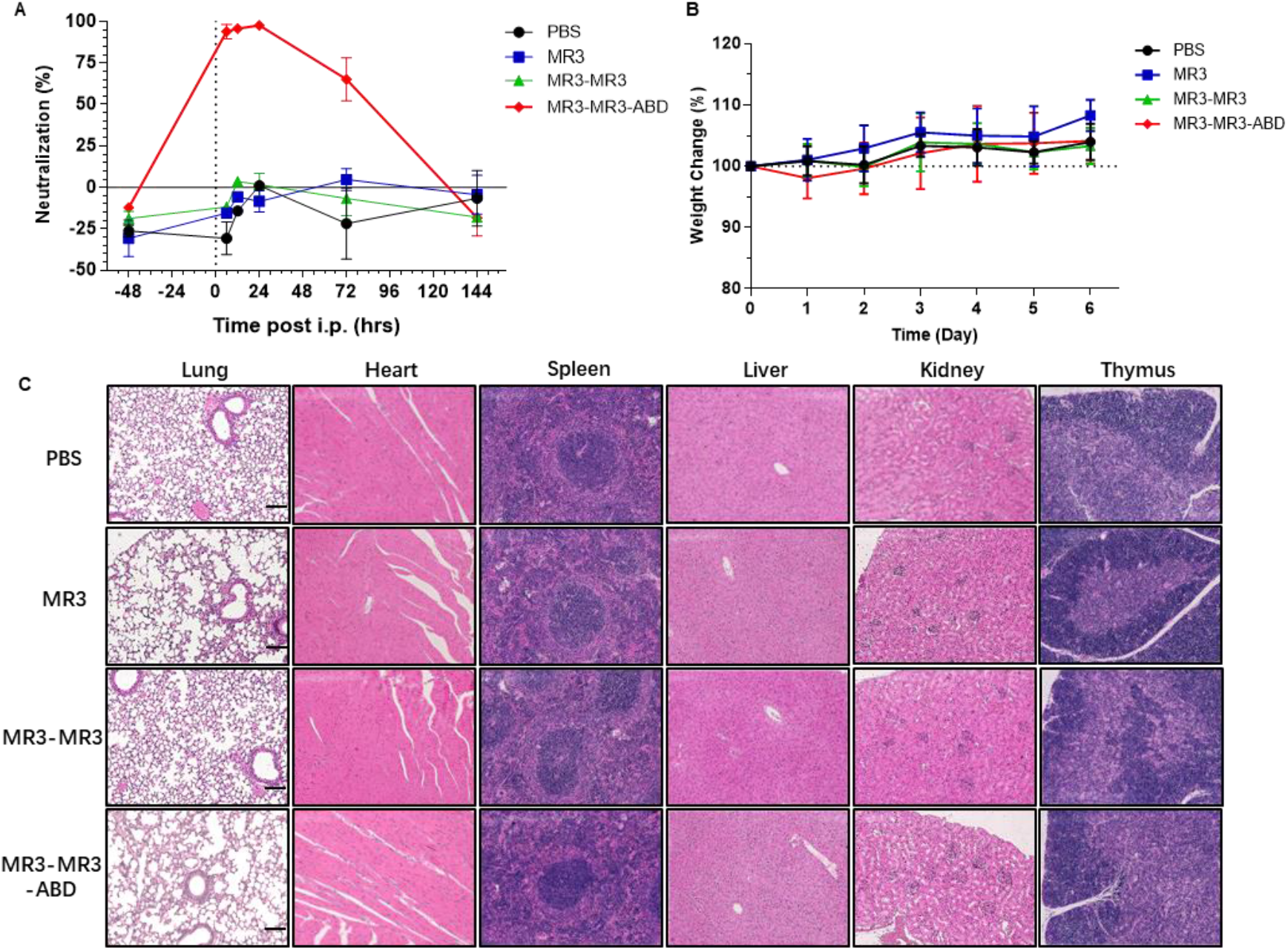
Evaluation of *in vivo* stability and toxicity of nanobodies. (**A**). Neutralization activity of sera from mice injected with sybodies. Sera were collected from mice injected with sybodies MR3, MR3-MR3, MR3-MR3-ABD, or PBS at the indicated time points. For neutralization assay, sera were preincubated with SARS-CoV-2 pseudovirus for 1 h before infection at 1/200 dilution. The infection rates on VeroE6-hACE2 were measure by FACS 3 days post infection. (**B**) Body weight changes. The body weight data are presented as means ± the SD of mice in each group (n= 4). No significant differences are observed. (**C**) Representative histopathology of the lungs, heart, liver, spleen, lungs, kidney, and thymus for the different sybodies injected. At day 3, the organ were collected, fixed, sliced and stain with hematoxylin and eosin. The images and areas of interest are magnified 100 ×. Bars indicate 100 μm.

